# Molpylot: a shared workspace for human–AI collaboration in molecular simulation

**DOI:** 10.64898/2026.11.24.768932

**Authors:** Stefano Motta, Anna Ranaudo, Lara Callea, Gianluca Santini

## Abstract

Rigorous biomolecular simulation requires a long chain of physical decisions across fragmented tools. Language-model agents offer to automate this work, yet current agents operate outside the scientist’s environment, generating ad hoc scripts or files that are effortless to launch but laborious to inspect and verify. Here we introduce Molpylot, an interactive modeling platform that unifies structure preparation, molecular dynamics, specialist protocols and replica-aware analysis in a graphical workspace designed to be immediate for the scientist, while exposing that same live session to any language model through 256 typed, tested tools. Operating within this shared workspace, the agent executes routine tasks under built-in safeguards that catch physical inconsistencies, while scientific judgment stays with the scientist. Across 18 systems, including asymmetric membranes, quantum-parameterized metal sites, enhanced sampling and free-energy calculations, we show how agents and built-in checks streamline setup, quality control and trajectory analysis. Molpylot is free for noncommercial use.

## Introduction

Molecular dynamics (MD) simulations are foundational tools for elucidating biomolecular mechanisms, ligand binding, and allosteric transitions. Yet executing a rigorous simulation demands a long chain of expert decisions: assigning protonation states, parameterizing cofactors and non-standard ligands, assembling membrane environments, setting up equilibration protocols, choosing collective variables, and enforcing statistical convergence across independent replicas^1,2^. In current practice, this workflow is fragmented across set-up servers, force fields, simulation engines, and analysis libraries, joined by ad hoc scripts whose assumptions are rarely documented, a persistent barrier to reproducibility and FAIR data stewardship^3^.

Large language models and autonomous agents offer an appealing route to lighten this burden^4,5^. Domain-specific agents now set up and run simulations from natural-language prompts^6–9^. However, most operate headless (running in background), and many have the model generate and execute its own scripts. Recent benchmarks expose the fragility of this paradigm: the best coding agents fully solve only 21%, 7%, and 4% of easy, medium, and hard MD tasks^10^ and complete fewer than half of realistic biomolecular workflows^11^. Crucially, their failures include physically incoherent set-ups (mis-specified force fields, inconsistent topologies, undersized membranes) that run without complaint^10,12^. Grounding agents in curated, expert-designed tools appears to mitigate this fragility: one MD agent completed 72% of tasks with such tools, against 28% for the same model given only a Python interpreter in which to write and run its own code^6^, in line with the broader case for tool-grounded autonomy in simulation-driven discovery^13^.

These failures exemplify a central dilemma of automation^14^: delegating pipelines entirely to headless agents produces executions too large to audit^15^ and invites automation complacency and illusions of understanding^16^, while justified distrust keeps AI out of core workflows. Given the growing recognition that generative AI lowers the cost of producing code without lowering the expertise needed to verify it^17^, full automation cannot be considered a safe option. Consequently, it is necessary to move to human-centered AI^18^ and mixed-initiative interaction^19,20^, systems that couple human agency and automated assistance in a shared workspace with explicit oversight.

Existing efforts to bring language models to molecular simulation fall into three families, each solving one problem while leaving the others open. Headless agents, whether they write code or call curated tools, accept a prompt and return files; the scientist sees the work only once it is finished^6,8,9,21^. Conversational agents attached to molecular viewers share visual state with the user, but their scope is structural inspection and analysis, they fall back on raw commands or generated Python for anything else, and none describes a simulation^22–25^. Commercial assistants in modeling suites (Schrödinger’s Maestro Assistant and Bunsen^26,27^, Cresset’s Flare AI Client^28^, and Nanome’s MARA^29^) aim for broader co-scientific capability, yet each is tied to one platform and to vendor-approved model providers, and none documents physical checks. Meanwhile, the Model Context Protocol (MCP)^30^ has emerged as an open standard for connecting models to tools across bioinformatics^31,32^, but MCP servers for simulation engines remain headless^33^.

None of these approaches provides what is actually needed: a live, synchronized session in which a scientist and any language model act on the same systems through identical, physically checked and reversible tools. Here we introduce Molpylot, a biomolecular modeling platform built around this shared workspace concept (Supplementary Video 1). By exposing its tools through MCP, with built-in physical checks and automated recording of every step and parameter, Molpylot makes human–AI collaboration both inspectable and reproducible. We then put it to work on a benchmark of 18 heterogeneous systems using only Molpylot’s MCP tools. Agents prepared and parameterized the simulation setups, deferring to the scientist whenever physical modeling choices were ambiguous, before auditing convergence and extracting key structural and thermodynamic properties. Molpylot is free for academic and other noncommercial use.

## Results

### A workspace in which the scientist can act directly as easily as delegating to the agent

A shared workspace between a scientist and a language model only works if the scientist can inspect, steer and verify the work as effortlessly as the model can execute it: whenever examining a structure, configuring a protocol or checking an observable takes more effort than delegating it, scientists are pushed toward unsupervised delegation^14^. At the same time, the workspace must span the entire modeling pipeline, because any missing step forces the agent to fall back on external scripts and uninspected files. Molpylot was built around these two design requirements: keeping every operation within immediate reach in the graphical interface while covering the modeling path from a database entry to a free energy calculation (Methods, Extended Data Fig. 1, Supplementary Table 1 and Supplementary Data 1). Within a single environment, the platform spans structure preparation, custom parameterization and simulation across biomolecular and drug-discovery workflows. Its capabilities range from sequence analysis, loop modeling and environment-aware protonation to bonded metalsite parameterization, asymmetric bilayer assembly, enhanced sampling, relative binding free-energy (RBFE) calculations, molecular docking and ligand-based QSAR.

To make verification equally immediate once calculations finish, 47 built-in analysis types (spanning trajectory measures, pocket geometry, membrane properties, allosteric networks and docking scores) feed into an interactive plotting environment linked to the 3D view (Supplementary Video 2). Independent trajectories loaded into an entry, or separate entries grouped together, are treated natively as replicas^2^, without duplicating coordinates: plots display individual replica traces, marginal distributions or replica means and standard deviations, while equilibration times^34^ and statistical uncertainties from autocorrelation and block averaging^35,36^ can be evaluated on demand. Analyses can be combined directly on screen: any per-frame plot can be colored by a second quantity or by conformational state, per-residue profiles can overlay an additional metric on a right-hand axis, and any two per-frame quantities can be projected onto a 2D free-energy surface. Clicking a point, a free-energy bin or a correlation-matrix cell jumps the 3D viewport to that frame or residue pair; conversely, dragging a rectangle on a plot lets the scientist extract those frames as a zero-copy sub-trajectory without duplicating coordinates in RAM, or assign them a named conformational state that can then be selected to color other plots of the entry (Supplementary Video 3). An integrated plot editor customizes layout, axes and series visibility and exports vector or high-resolution figures alongside complete CSV tables. Key preparation, simulation, docking and free-energy components were validated against reference engines and experimental benchmarks (Methods and Extended Data Table 1).

### Agents and scientists share a workspace

Molpylot is organized around a single stateful session (holding entries, trajectories, selections, 3D views, analysis results and history) operated through a unified catalog of 256 typed tools across 32 functional areas, exposed identically to the graphical interface, the built-in Python console and any MCP client through an embedded or stand-alone server^30^ (Fig. 1a; Supplementary Table 1).

**Fig. 1.**
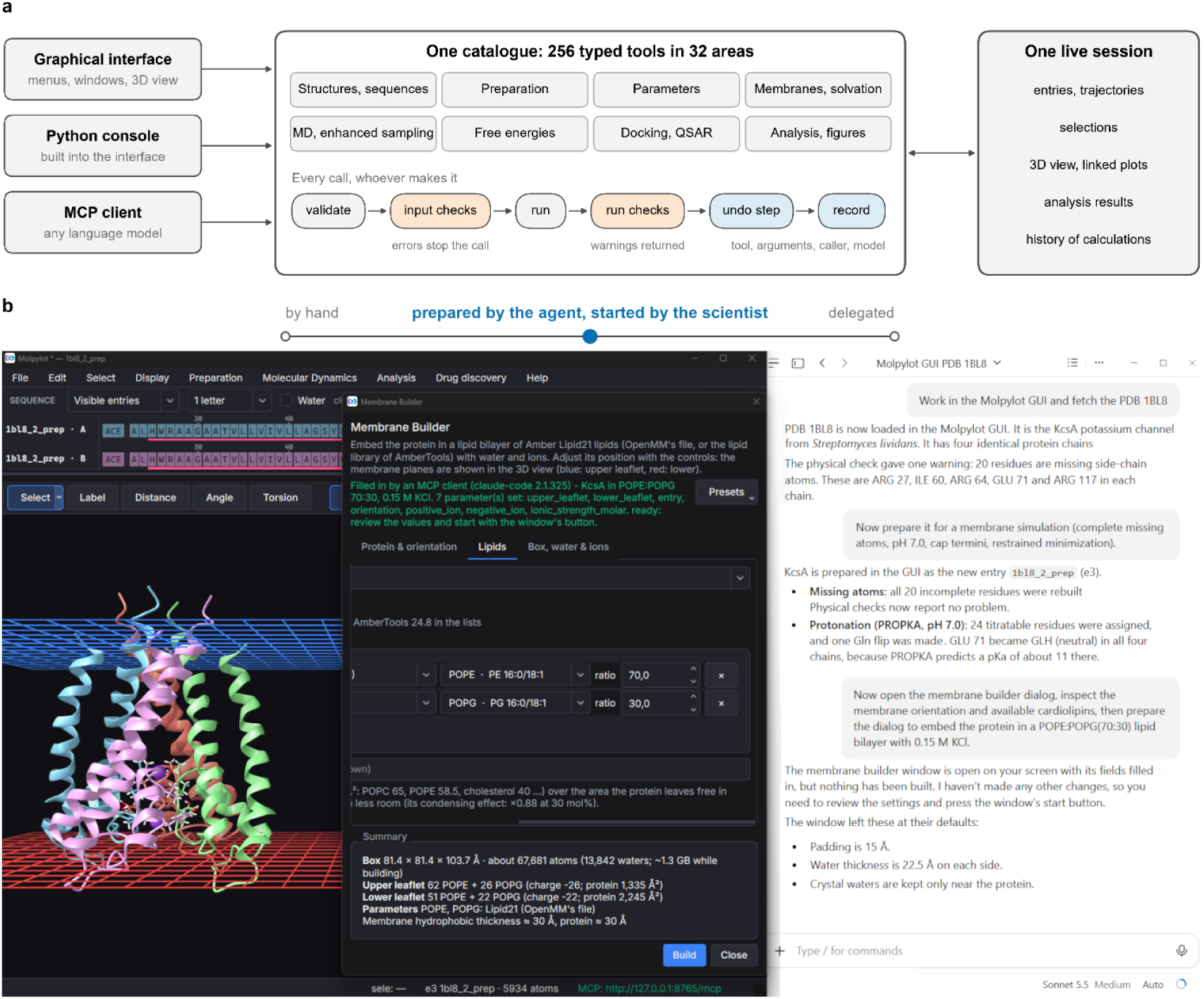
Architecture and live shared workspace of Molpylot. a, The graphical interface, the built-in Python console and MCP clients invoke the same 256 typed tools across 32 areas. Every call, whether initiated by the scientist or by a language model, undergoes argument validation and, where they apply, physical input and run checks, undo registration and provenance logging on a single live session. **b,** Delegation as a continuous dial during the preparation of the KcsA potassium channel (PDB 1BL8). In the client conversation (right), the agent fetches the crystal structure, reports the physical-check warning on 20 incomplete residues, rebuilds and protonates the tetramer, and pre-fills the Membrane Builder module for a 70:30 POPE:POPG bilayer with 0.15 M KCl. In the Molpylot window (left), the prepared structure is shown oriented between the upper (blue) and lower (red) membrane planes in the 3D viewport, with the pre-filled dialog open and flagged in green for the scientist to review and launch.

Because the graphical interface and the agent call the same underlying functions, every change to the session (humanor AI-initiated) is subject to the same physical checks and activity logging and can be undone in a single step or rolled back if interrupted. Any change made by the agent appears immediately in the 3D view and plots, and any selection or adjustment made by the scientist is instantly available to the agent. Scientists and agents do not exchange scripts or files; they share a workspace. This symmetry turns delegation into a continuous, step-by-step choice (Fig. 1b). A scientist can inspect a structure, ask an agent to parameterize a ligand and assemble a bilayer, adjust the orientation in 3D, and hand back equilibration and analysis, with every step visible and reversible. Between manual operation and autonomous execution sits an intermediate mode (prefill_module) in which the agent populates and validates the fields of a graphical dialog, leaving the scientist to review the settings and start the calculation. Agents operate across the platform without custom fine-tuning or loading the manual into the prompt. Instead, they query integrated discovery and local documentation tools to retrieve relevant parameters, defaults, selection grammar and procedural recipes on demand (Methods).

Rather than executing model-written code, each tool runs a tested protocol, and the tools are wrapped by built-in physical safeguards (Methods, Supplementary Table 2 and Supplementary Data 2). Before execution, these rules check for physical inconsistencies such as severe steric clashes, chain breaks, unparameterized ligands or non-neutral boxes; following MD simulations, they audit trajectories and logs for thermodynamic drift, lost metal coordination or poor sampling overlap. Critical errors stop a call before it runs, whereas warnings and suggested remedies are returned directly to both the agent and the scientist, appearing in the chat transcript or, for actions started in the graphical interface, in the interface itself.

Finally, provenance is recorded across all calculation tools and background analyses while automatically writing a complete Materials and Methods section with software versions and DOI citations into self-contained job folders for cluster execution.

### Agents confined to the tools prepared and analyzed eighteen systems

To test Molpylot’s protocols and shared workspace, we ran an 18-system validation campaign spanning soluble proteins, protein–ligand complexes, lipid bilayers, membrane receptors, covalent and phosphorylated complexes, metalloproteins, a conformational landscape and RBFE networks (Fig. 2a; Extended Data Table 2). Both phases used language-model agents restricted to Molpylot’s MCP tools (Methods). First, preparation agents assembled pure lipid bilayers from target compositions or took PDB entries through repair, protonation, quantum-chemical parameterization and membrane embedding, solvating every system into a ready-to-run simulation package. Second, analysis agents audited convergence and extracted properties from the produced trajectories (Supplementary Sections 1–5 and Supplementary Note 2).

**Fig. 2.**
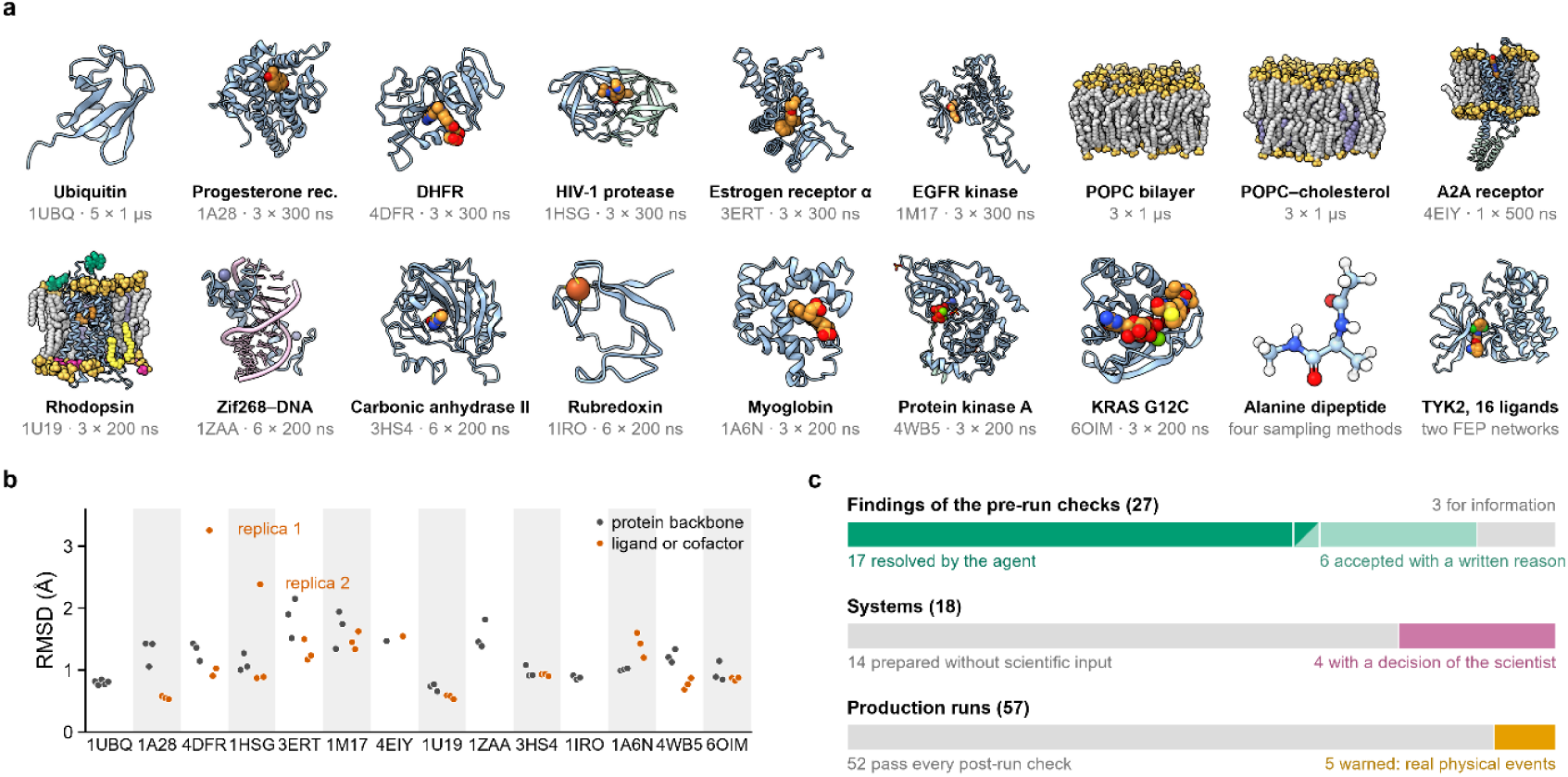
Agents confined to the tools prepared and analyzed 18 systems. **a**, The 18 systems, with the number and length of the production runs; the three metalloproteins with a bonded and a non-bonded model have three runs of each. Proteins are shown as prepared, the two bilayers and rhodopsin at the end of a run, and the A2A receptor after equilibration; water and bulk ions are not drawn. **b**, Mean RMSD from the prepared structure in the analyzed window of every replica: protein backbone (gray; the folded core or the transmembrane helices as indicated in Supplementary Table 5) and ligand or cofactor (orange). The two labeled replicas are events that the agents reported for a single replica: the glutamate tail of methotrexate in PDB 4DFR and the pyridylmethyl group of indinavir in PDB 1HSG. **c**, Summary of check outcomes and human interventions across the benchmark, showing how the 27 pre-run check findings were handled (top), how many of the 18 systems required a human decision (middle), and how many of the 57 production trajectories passed or triggered warnings in post-run checks (bottom).

Across the benchmark, folds, bilayer structures, cofactor sites and ligand poses were maintained (Fig. 2b; Extended Data Fig. 2), with the agents reporting local rearrangements such as a water replacing an ATP oxygen in the Mg²⁺ shell of protein kinase A (Extended Data Fig. 2d–f; Supplementary Table 5). The logs show a clear division of labor between agent, checks and scientist (Fig. 2c; Supplementary Table 4):

- **Routine defects resolved by the agent.** Pre-execution checks returned 27 distinct findings, none of which stopped a call. The agent resolved most warnings itself, including repairs of missing heavy atoms, unmodeled loops, clashing waters and unparameterized ligands; accepted 6 with a written reason (such as a high sulfonyl-sulfur charge and the non-bonded metal controls); and kept the *cis* peptides of dihydrofolate reductase flagged for information.
- **Scientific ambiguities passed to the scientist.** During initial screening, the agent paused on an additional candidate, streptavidin (PDB 1STP), whose PDB entry contains only one subunit of the functional tetramer; because no tool reconstructs symmetryrelated assemblies, we excluded it from the final benchmark. Of the 18 systems, 14 needed no scientific input; on the other four, the agent flagged crystals with two copies in the asymmetric unit (PDB 4DFR and 1A28), which we reduced to one chain, and asked about the distal His64 tautomer of myoglobin (PDB 1A6N), where we chose HIE, the tautomer that neutron diffraction finds hydrogen-bonded to the bound oxygen in oxymyoglobin^37^; and in KRAS G12C (PDB 6OIM), the agent kept the default Mg²⁺ model and offered the 12-6-4 model only as an option, which the scientist then chose as the model that better describes Mg²⁺ bound to phosphates.
- **Post-run warnings and replica-specific events.** Post-run checks passed 52 of 57 trajectories and warned on 5, each a real physical event rather than a setup error: the C-terminal tail of EGFR (PDB 1M17) drifting from its crystal position and, in two replicas, reaching its periodic image, and the asymmetric rhodopsin bilayer (PDB 1U19) remaining incompletely equilibrated in lateral area across two replicas (Supplementary Fig. 1k; Extended Data Fig. 3b; Supplementary Table 5). Beyond the automated warnings, the analysis agents reported single-replica conformational events rather than averaging them away: a ubiquitin loop sub-state, a rearranged methotrexate glutamate tail in PDB 4DFR, and the indinavir pyridylmethyl group leaving its subsite and returning in PDB 1HSG (Supplementary Fig. 1; Supplementary Table 5 and Supplementary Data 3).

We next examine how this collaborative workflow handles four non-routine applications where setup, parameterization or sampling choices are critical.

### Agent and scientist working together on four demanding cases

Four areas of biomolecular simulation (building post-translationally modified membrane proteins into asymmetric bilayers, parameterizing metal sites from quantum chemistry, comparing enhanced-sampling estimators and running RBFE campaigns) lie outside the routine molecular-dynamics toolkit and usually require specialist expertise. These four cases illustrate what Molpylot’s tools make accessible within one workspace, and how the scientific work divides between agent and scientist.

### Asymmetric receptor membranes from a biological brief

Molpylot builds symmetric or asymmetric bilayers from a lipid composition, orients membrane proteins within them, and during analysis evaluates lipid area convergence and calculates properties by leaflet and lipid species (see Supplementary Fig. 2, Supplementary Table 5 and Supplementary Data 4 for pure POPC, POPC–cholesterol and the A2A adenosine receptor). As a demanding test case, we prepared dark-state bovine rhodopsin (PDB 1U19), which combines in a single system a covalent 11-*cis* retinal on a protonated Schiff base, two intradiscal *N*-glycans, two cytoplasmic palmitoyl anchors, buried carboxylates with non-standard protonation states, and a native rod-disc membrane whose two leaflets differ in headgroup charge, polyunsaturated acyl chains and cholesterol content. To assemble this system, the agent paired compatible protein, lipid, glycan and covalent-residue force fields, assigned the dark-state protonation of Asp83 and Glu122 from spectroscopic data, and chose the acyl chains of seven lipid species. When it found a mis-modeled mannose in the Asn15 glycan, it deleted the sugar and opened its report by asking the scientist to review that judgment call; rather than trusting the automatic orientation, it also verified landmark positions from the built coordinates to confirm that “Each composition is on the correct side”. Over 3 × 200 ns (Fig. 3a–c; Extended Data Fig. 3a–d; Supplementary Table 5), the receptor remained inactive, the retinal stayed 11-*cis*, and the bilayer preserved its asymmetry with all phosphatidylserines retained in the cytoplasmic leaflet.

**Fig. 3.**
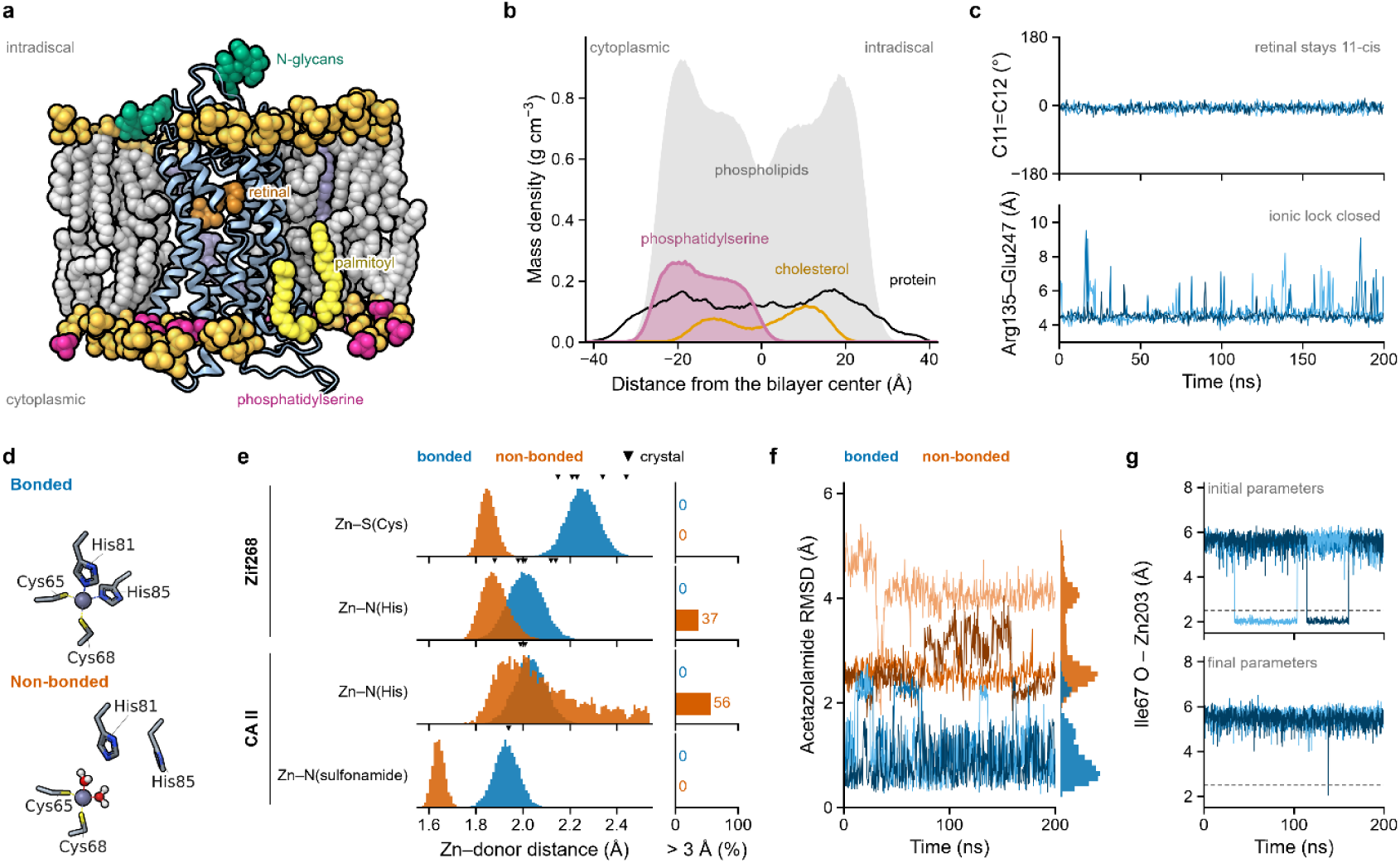
A receptor in an asymmetric membrane and two zinc sites. **a**, Rhodopsin (PDB 1U19) in the disc membrane that the agent built from the biological brief, after 200 ns (replica 1), showing the 11-*cis* retinal, the two *N*-glycans on the intradiscal side, the two palmitoyl cysteines, and the phosphate groups of the lipids; those of phosphatidylserine (magenta) are all in the cytoplasmic leaflet. Lipid tails are drawn only behind the receptor. **b**, Mass density across the membrane in the analyzed windows of the three replicas of 200 ns: all phospholipids (gray), phosphatidylserine, cholesterol and the protein. **c**, The C10–C11=C12–C13 torsion of the retinal and the Arg135–Glu247 distance (the ionic lock) in the three replicas. **d**, The Zn203 site of Zif268 in replica 3 at 121 ns of the bonded and of the non-bonded model; in the latter both histidines have left the zinc and two waters have taken their place (dashed lines, coordination). **e**, Zinc–donor distances in the analyzed windows, bonded against non-bonded (3 × 200 ns per model), for Zif268 and carbonic anhydrase II; triangles, distances in the crystals; right, the share of frames with the donor beyond 3 Å. **f**, Heavy-atom RMSD of acetazolamide in carbonic anhydrase II over the whole production of the three replicas of each model, with its distribution. **g**, Distance of the carbonyl oxygen of Ile67 from Zn203 in the bonded runs of Zif268: with the initial parameters the run check reported it within 2.5 Å (dashed line) in 35% and 23% of the frames of two replicas; with the final parameters it comes that close for 0.5 ns in one replica.

### Quantum-chemically parameterized metal sites

For metalloproteins the agent, using Molpylot tools, identifies the coordinating donors, runs the quantum-chemical calculations and builds a bonded model (with Seminario force constants and RESP charges from B3LYP/631G* calculations^38–40^; Supplementary Table 6) or retains the force field’s non-bonded ion as a control. Over 3 × 200 ns per model (Fig. 3d–f; Supplementary Fig. 3; Supplementary Table 5), the quantum-derived bonded sites of Zif268 and carbonic anhydrase II, like the Fe(III) site of rubredoxin (Extended Data Fig. 4), preserve their tetrahedral coordination in every replica, with no water entering the first shell and the acetazolamide inhibitor of carbonic anhydrase II remaining locked in its crystal pose. By contrast, the standard non-bonded zinc model collapses in both systems: in Zif268 it over-shortens the Zn–S bonds^41^ and loses half of its histidine coordination interactions across the replicas, while in carbonic anhydrase II it loses a coordinating histidine during equilibration and recruits neighboring protein and ligand oxygens instead, displacing acetazolamide from its binding pose. These two systems also illustrate how the agent and the built-in checks catch subtle coordination artifacts during both preparation and analysis. During preparation of carbonic anhydrase II, the agent inspected the active site after energy minimization: seeing that an initial minimization pulled a sulfonamide oxygen onto the zinc, it rejected the distorted geometry (”That would have made a false fifth donor”), repeated the preparation without heavy-atom minimization, and flagged that protocol deviation for the scientist before the jobs were submitted (confirming in the final setup that the crystallographic distance was preserved). On Zif268, after an initial bonded parameterization with HF/6-31G* RESP charges, the post-run check reported unprompted a non-donor atom at one zinc, “held by no bonded term: ILE 67/C O within 2.5 Å in 35% of the frames”. Directed by the scientist and executed by the agents, the sites were parameterized again, with B3LYP/6-31G* RESP charges and a revised bonded model, and the contact now lasted only 0.5 ns in one replica of three (Fig. 3g).

### Enhanced sampling across four protocols

Setting up and comparing distinct enhancedsampling estimators normally requires separate inputs and post-processing workflows. To test them within one workspace, we used alanine dipeptide (a standard model peptide whose conformations are described by the two backbone angles φ and ψ) to compute the free-energy difference and the transition barrier between the two halves of its Ramachandran map. An agent configured plain MD, well-tempered metadynamics^42^, umbrella sampling with WHAM^43^ and steered dynamics with Jarzynski’s equality^44^ from Molpylot’s defaults, and the analysis agent compared them in a single project using basin boundaries fixed once for all methods from the metadynamics barriers (Fig. 4; Supplementary Table 5). All four methods agree on both the free-energy difference between the two regions and the height of the barrier near φ = 0° (Fig. 4a,b), while the analysis agent evaluated the convergence and limitations of each estimate (Fig. 4c; Extended Data Fig. 3e–h; Supplementary Section 5). The agent also warned that 500 ns of plain dynamics were not yet tightly converged (”Only at the ±0.25 kcal/mol level”), whereas metadynamics converged after about 40 ns. In umbrella sampling, neighboring windows overlapped well, yet the agent noticed that the initial segment of each window gave a lower estimate than the remainder (”a systematic transient at the start of each window, not chance”); it reported the result with and without that segment (”Choosing between them is up to the author”), and the scientist excluded it, with the result insensitive to the exact cut. For steered dynamics, the agent predicted from the restraint stiffness a barrier slightly too low, within the bootstrap uncertainty of the profile, consistent with its close agreement with umbrella sampling.

**Fig. 4.**
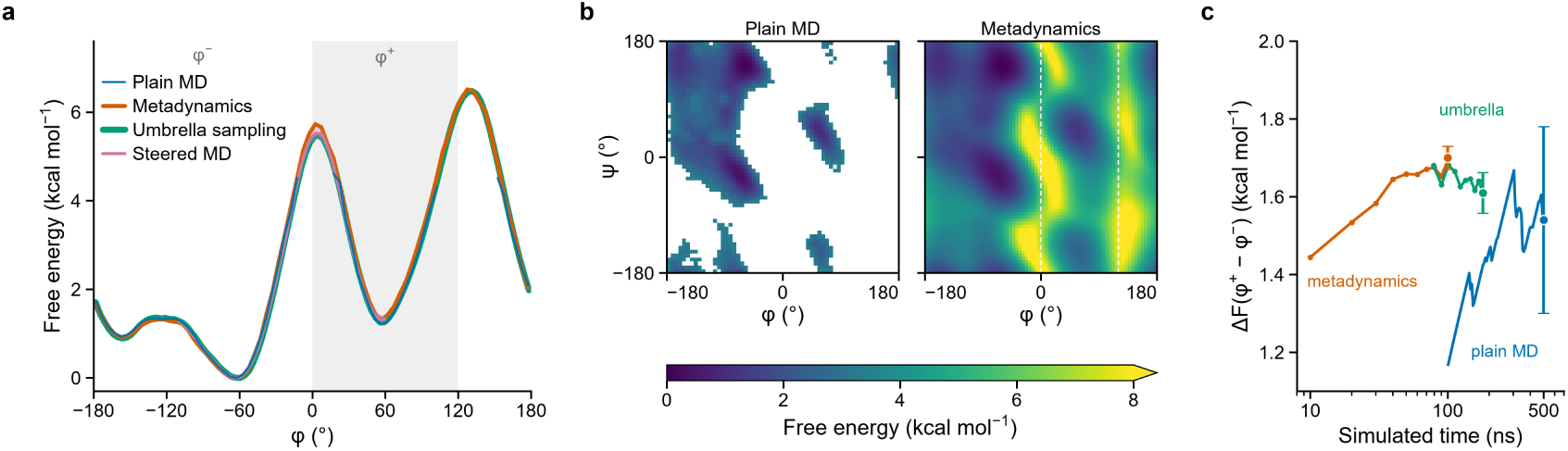
Four sampling protocols on one free-energy profile. **a**, Free energy of alanine dipeptide along φ from 500 ns of plain MD (drawn where sampled), 100 ns of well-tempered metadynamics, umbrella sampling (36 windows of 5 ns, the first 1.25 ns of each left out, WHAM) and steered dynamics (20 pulls from −110° to +60°, Jarzynski’s equality; shading, bootstrap error). Every profile is zero at its lowest point for φ < 0°; the band is the φ⁺ basin (0° ≤ φ < 120°). **b**, Ramachandran free-energy surfaces from plain dynamics and from metadynamics on one color scale (white, no samples; dashed lines, φ = 0° and 120°). **c**, The free-energy difference between the two basins against the simulated time it uses: the cumulative estimate of plain dynamics from 100 ns (before, it rests on the first few crossings and ranges from 0.98 to 4.4 kcal mol⁻¹), metadynamics after every 10 ns, and umbrella sampling with every window used from 1.25 ns up to 2.2–5 ns (79–180 ns in all). Points with bars, the final estimates and their errors (1.54 ± 0.24, 1.70 ± 0.03 and 1.61 ± 0.05 kcal mol⁻¹).

### Relative binding free energies on an agent-designed network

Molpylot helps plan, map, prepare and analyze RBFE campaigns with the OpenFE protocol, while the scientist supervises the network and every perturbation edge in the graphical interface (Fig. 5a). For the 16 ligands of the TYK2 benchmark series^45^, one agent set up the published 22-edge network, whereas a second received only the receptor, the posed ligands under anonymous names, a budget of about 1,000 GPU-hours and the instruction not to look the compounds up, so that it could not reuse the published network. The second agent planned a minimally redundant network of 30 edges, pruned it to 24 and chose three repeats per leg. Against experimental data, the published network gives an RMSE of 0.60 kcal mol⁻¹ and the agent’s network 0.59 kcal mol⁻¹, with similar correlation and ranking statistics, close to published OpenFE results on these ligands^46^ (Fig. 5b; Supplementary Table 7 and Supplementary Data 5–7). The analysis agent then separated sources of error that a single RMSE obscures (Fig. 5c). The same ligands, ejm_49 and ejm_55, are the worst outliers in both networks, as in the published results, pointing to an error due to the force field or to the binding mode rather than in the network or sampling, whereas edges with large repeat-to-repeat spread were flagged so that the scientist could extend their simulations.

**Fig. 5.**
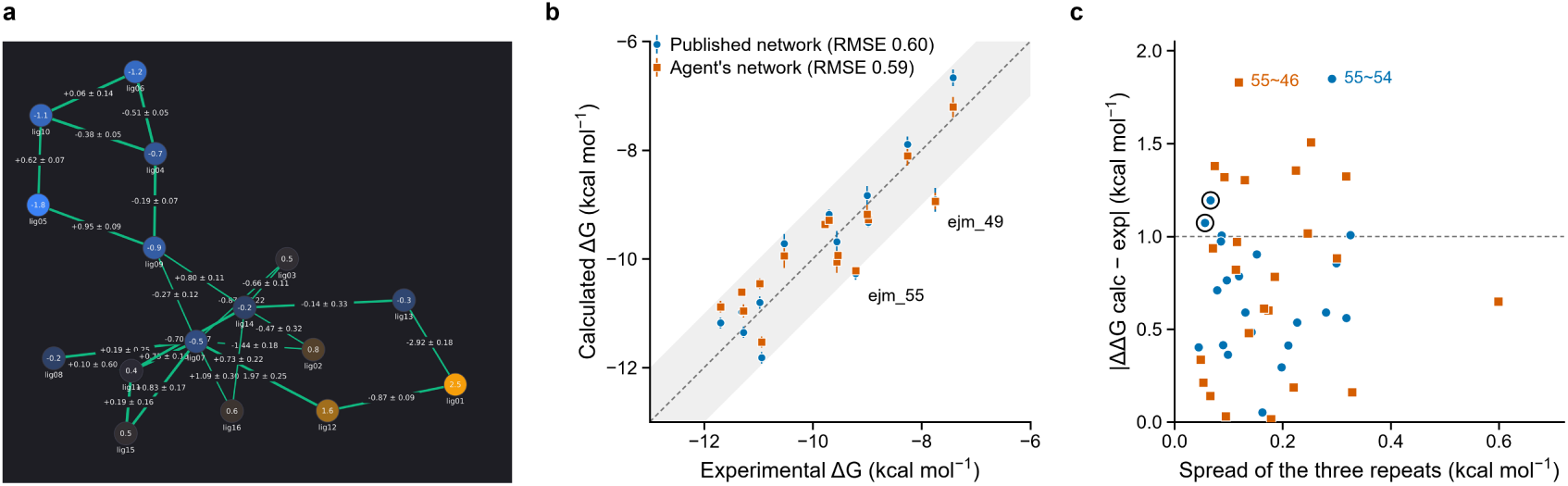
An agent-designed free-energy network against the published one. **a**, The agent’s campaign in the graphical interface: its network of 24 edges over the 16 ligands of TYK2, with the calculated ΔΔG and the spread of the repeats on every edge. **b**, Binding free energies (ΔG) calculated with the published network (22 edges) and with the agent’s, against the measured ones; bars, spread of the three repeats; band, ±1 kcal mol⁻¹. The ligands ejm_49 and ejm_55 are the worst outliers of both. **c**, Absolute error of every edge against the spread of its three repeats. Ringed, the two edges of the published network whose errors of 1.07–1.19 kcal mol⁻¹ come with spreads of 0.06–0.07 kcal mol⁻¹; labeled, the worst edge of each network, both through ejm_55.

## Discussion

Computational molecular science relies on complex, multi-stage workflows that traditionally force researchers to chain together fragmented command-line tools and scripts. Languagemodel agents promise to automate this work, yet most operate outside the scientist’s environment, generating headless pipelines or ad hoc code that is effortless to launch but laborious to verify^17^, so that errors may pass undetected into scientific software^47^. Whenever verifying an automated action takes more effort than delegating it, human oversight is at risk^14^. Molpylot addresses this challenge by redesigning the modeling platform itself: first bringing the entire simulation and analysis stack into an interactive graphical workspace designed to be immediate for the human scientist, and then placing the language model inside that same live session. Because human and model act on the same workspace through the same tools, delegation becomes a step-by-step supervised choice rather than an all-or-nothing handover^20,48^.

Within this shared workspace, the software, the agent and the scientist play distinct, complementary roles. Built-in physical checks enforce self-consistency before calculations start and audit thermodynamic and structural stability once they finish, catching setup defects or trajectory anomalies. The agent handles the operational mechanics (chaining tools together, configuring protocols through built-in documentation and synthesizing quantitative diagnostics across replicas) without ever writing uninspected code. Choosing physical approximations, interpreting unexpected results and judging sampling adequacy remain with the scientist, who can inspect and adjust any agent action directly in the 3D viewport and linked plots. Across 18 systems, agents carried both routine and specialist workflows largely end to end, with the scientist stepping in when needed. The resulting simulations preserved crystallographic geometries and agreed with experimental benchmarks.

This design sets Molpylot apart from existing approaches to AI in molecular modeling. Headless agents sacrifice interactive oversight^6,8,9,21^; conversational agents attached to molecular viewers lack a simulation and free-energy engine^22–25^; and commercial assistants are converging on parts of this design^26–29^. To our knowledge, no other published system combines an end-to-end simulation stack, a live shared session, physical safeguards and provenance in the tools, and a free choice of model. This model independence also gives the platform long-term durability: because Molpylot connects to agents through an open protocol, the underlying language model can be changed without altering the tools, the checks or the scientific record as providers revise and retire hosted models^49,50^. The deliberate price of this architecture is that an agent can only execute protocols present in the tool catalog (new methods require reviewed developer code^51^), while external client connections introduce prompt-injection surfaces (mitigated as detailed in Methods^52,53^) and rely on client-declared model identifiers for provenance.

Provenance, however, records what was done and at whose request, not who understood it^17^. In scientific training, delegating early to AI agents risks accruing epistemic debt^54^, leaving junior researchers able to produce complex simulations without developing the physical intuition needed to evaluate them^17^. Because manual execution in Molpylot remains as direct as delegation, trainees can first master the methods by hand and only later add gradual, supervised assistance within the same environment.

More broadly, Molpylot offers a different blueprint for scientific software in the era of AI. Rather than treating language models as autonomous script writers that bypass the user interface, software can be built so that human and model share the same workbench, the same safeguards and the same record. Molpylot applies this design to biomolecular simulation, giving researchers an environment with built-in physical checks in which language-model assistance expands what they can tackle, while every result remains one they can inspect and defend.

## Methods

### Software architecture and session model

Molpylot is written in Python 3.11 around a central stateful session built on NumPy and SciPy^55,56^ for vectorized coordinate and topology storage, OpenMM 8.6.1^57^ for molecular mechanics and simulation execution, ParmEd^58^ for reading, editing and writing Amber and GROMACS topologies, and MDAnalysis 2.11.0^59^ for trajectory I/O and analysis. Domainspecific engines for structure preparation, small-molecule chemistry, quantum mechanics, freeenergy calculations and docking are integrated directly or managed in isolated environments by a built-in component manager.

At runtime, the session maintains molecular entries (each bundling atomic coordinates, topology, trajectories, visual representations, force-field parameters, analysis outputs and provenance records) together with named atom selections in thread-safe registries, publishing state updates to the PySide6 graphical interface through a batched event bus. To keep memory overhead low when inspecting multi-replica datasets, trajectory sub-ensembles (such as conformational clusters or regions brushed on a plot) are stored as zero-copy frame-index views; coordinate superpositions and periodic-box imaging are applied on the fly as frames are read, and independent replica trajectories are grouped without duplicating coordinates on disk or in memory.

Every tool call initiated from the graphical interface or a connected MCP client executes on a worker thread under a unified task controller that streams progress updates and supports clean cancellation (terminating external subprocesses and stopping OpenMM minimizations or quantum-chemical SCF cycles at the next checkpoint). Each session-modifying tool call forms a single step in a reference-based undo history (40 steps by default), and interrupted or failed calls automatically roll back partial state changes. Project files serialize the session state without Python pickle, ensuring that opening a saved project never executes arbitrary code.

### Rendering

The interactive 3D viewport is built on ModernGL using the OpenGL 3.3 core profile. Atomic positions and colors reside in GPU textures, while geometric representations (ray-cast sphere impostors, instanced bond cylinders and Catmull–Rom backbone cartoons swept on the GPU) rebuild their vertex buffers only when representations change, so trajectory playback updates only the coordinate texture. Solvent-excluded molecular surfaces are triangulated on a background thread from a probe-accessible distance field via surface nets and anchored to moving atoms during playback with periodic exact recomputation. For publication figures, Blinn–Phong lighting and depth cueing are augmented with screen-space and per-atom ambient occlusion and depth-buffer silhouettes. The same rendering pipeline runs headlessly through EGL on a GPU or software rasterizer, allowing the capture_viewport tool to return live 3D images to language-model agents.

### MCP server, tool layer and clients

Molpylot exposes 256 shared catalog tools across 32 functional areas, plus four MCP-specific discovery and job-management tools (find_tools, get_job, list_jobs and cancel_job), through a server built on the MCP Python SDK 2.3.0 (Supplementary Table 1 and Supplementary Data 1). Clients connect either to the server embedded in a running GUI window (over loopback streamable HTTP, acting directly on the scientist’s live visual session) or to the stand-alone molpylot-mcp server (over stdio, SSE or streamable HTTP). Session-modifying calls run sequentially inside an undo and provenance scope, streaming progress notifications at up to 1 Hz; operations exceeding 40 s (configurable) return a background job identifier that the agent tracks asynchronously. When a GUI server is active on the same machine, headless server responses include a notice pointing to the live session’s address.

Because loading all MCP tool schemas at once would consume roughly 560,000 characters of model context, the server provides five built-in tool profiles (full, essential, simulation, analysis and drug design; Supplementary Table 3) as well as custom user-defined profiles. In the embedded server, switching profiles updates the active toolset in place and notifies connected clients via notifications/tools/list_changed without restarting the session or interrupting running jobs. Tool schemas remain compact by capping the opening summary of each description at 200 characters and omitting long parameter explanations. Instead, agents discover relevant tools through find_tools, a local TF-IDF router that ranks tools for a task description and returns multi-step workflow recipes without invoking a language model, and retrieve detailed parameter documentation on demand via search_docs, which queries 300–500-token chunks of a tool reference generated from the live catalog and of workflow guides using BM25 and optional dense embeddings. Six standard workflow recipes are also exposed as MCP prompts, and get_module_defaults paired with prefill_module allows an agent to populate a GUI dialog for the scientist to review and launch.

To bound execution risks, no tool runs model-generated code, and dedicated deletion tools carry the MCP destructiveHint annotation. Filesystem access is restricted to the project directory, a private temporary workspace and user-authorized paths; HTTP connections require a locally generated bearer token and reject requests carrying foreign Host or browser Origin headers. External text returned by tools (such as structure headers, database records, literature abstracts, BLAST descriptions or log files) is stripped of control, invisible and bidirectional characters, labeled as external data in an external_text field, and scanned with regular expressions for English instruction-like patterns to mitigate indirect prompt injection (Supplementary Note 1).

### Physical checks and provenance

Twenty-two tools that prepare structures, parameterize molecules, assemble systems, run or export simulations, and analyze trajectories or free energies are wrapped by automated physical safeguards (including pre-execution input rules on 11 tools), covering the 80 finding codes they expose, classified as error, warning or information (Extended Data Fig. 1a and Supplementary Table 2; every rule with its threshold and remedy in Supplementary Data 2). These checks are implemented in Molpylot itself, in addition to any checks built into the engines and libraries it calls. Before execution, input rules verify structural and topological consistency (catching defects such as missing heavy atoms, steric clashes, unparameterized ligands or incompatible integration time steps); after execution, output rules audit simulation logs, trajectories and free-energy results for thermodynamic drift, loss of metal coordination or poor sampling overlap. Under the default strict policy, error-level input findings block execution and job-folder export, whereas warnings and remediation suggestions are returned in the tool output; internal check exceptions are logged without halting the tool.

All 93 calculation tools record structured provenance on the affected entries and in the session log. Each record captures both the computational specification (the tool name, timestamp, non-default arguments, input/output files and Molpylot version) and the actor (whether the call came from the GUI, console, script or MCP client, whether prefill_module was used, and the model identifier from the server environment or the client’s unverified declaration). When writing input folders for simulations, docking or free-energy campaigns, a built-in methods generator compiles the relevant provenance chain into a past-tense Materials and Methods section complete with software versions and a BibTeX file drawn from 183 DOI-verified references.

### Structure preparation and parameterization

**Proteins and nucleic acids.** Structure preparation optionally removes waters and crystallization additives, converts selenomethionines, trims truncated termini, adds terminal caps, and reconstructs missing side chains and loops with PDBFixer^60^, falling back on cyclic coordinate descent^61^ for loops and rotamer libraries^62^ for side chains if geometric checks or steric overlaps fail. Missing nucleotide atoms are completed, residue protonation states are assigned with PROPKA 3^63^ in a two-pass scheme that accounts for the electrostatic environment of bound ligands, cofactors, glycans and metal thiolates, and hydrogen-bond networks are optimized with PDB2PQR^64^. Prepared structures are energy-minimized under heavy-atom restraints; during this step, unparameterized free ligands receive provisional OpenFF Sage 2.3.0 parameters with NAGL charges^65^, while incomplete ligand copies, metal clusters or covalent adducts are held fixed as charged UFF Lennard-Jones spheres.

### Small molecules, cofactors and metal sites

Using RDKit^66^ and the OpenFF toolkit^67,68^, ligand bond orders and formal charges are perceived from 3D geometry (bond angles, planarity and mixed-integer linear programming for Kekulé assignment), existing hydrogens, SMILES/SDF/MOL2 inputs or the PDB Chemical Component Dictionary, and protonation and tautomeric states are scored with an electrostatically coupled p*K* ₐ model. By default, small molecules are parameterized with OpenFF Sage 2.3.0 and NAGL graph-convolutional charges^65^. Alternatively, GAFF or GAFF2^69^ topologies can be generated through AmberTools^70^ and paired with two-stage Merz–Kollman RESP charges^39^ fitted to HF/6-31G* electrostatic potentials from PySCF^71^ and geomeTRIC^72^ (optionally multi-conformer and GPU-accelerated, with scanned halogen extra points and QM-refitted dihedral terms), AM1-BCC^73^, MMFF94 or Gasteiger charges. Post-translationally modified residues and covalent ligand adducts are parameterized as ACE/NME-capped fragments combining Amber backbone types with OpenFF Sage side-chain terms.

Metal centers follow the bonded MCPB protocol of Li and Merz^40^: after perceiving the coordination graph, Molpylot derives Seminario bond and angle force constants^38^ from a B3LYP/6-31G* Hessian of the small model and RESP charges from the capped large model (verifying SCF stability and Hessian eigenvalues) while setting torsions across the metal to zero; empirical force constants or the zinc ZAFF/EZAFF sets^74,75^ are available when quantum chemistry is not used. Supplementary Table 6 lists, for the validation sites, the donors, the two models, their net charges and spin multiplicities. Standard cofactors use curated Amber parameter sets: Meagher et al. for ATP, ADP, GTP and GDP^76^; Ryde and co-workers for NAD⁺, NADH, NADP⁺ and NADPH^77–79^; GAFF2 with RESP charges for oxidized FAD and FMN; the AmberTools heme models of Shahrokh et al. for cysteine-ligated heme^80^ and Giammona–Bayly for histidine-ligated or pentacoordinate Fe(II) heme; and GLYCAM-06j for covalently linked glycans^81^.

### Molecular dynamics and membranes

Soluble systems default to Amber ff19SB^82^ with OPC water^83^, whereas membrane systems default to ff14SB^84^ with TIP3P^85^ and Lipid21^86^; nucleic acids use OL21 or OL15 DNA^87,88^ and OL3 RNA^89^, with alternative water models^85,90,91^ and Joung–Cheatham or Li–Merz ion parameters^92,93^ supported. Systems are assembled via OpenMM templates and can be run directly in OpenMM or exported with ParmEd to Amber and GROMACS^94^ formats. Default dynamics settings use particle-mesh Ewald electrostatics (1.0 nm cutoff)^95^, constrained bonds to hydrogen with rigid water, a Langevin middle integrator^96^ at a 2 fs time step (4 fs with hydrogen-mass repartitioning ^97^) and a Monte Carlo barostat^98^ (semi-isotropic for membranes). Equilibration presets include the ten-stage Roe–Brooks protocol^99^ (repeating final NPT stages until density converges), a standard heating and restraint-release protocol, and a six-stage membrane relaxation scheme.

To embed membrane proteins, Molpylot determines the transmembrane orientation via a hydrophobic-belt search using the Hessa transfer-energy scale^100^, packs symmetric or asymmetric bilayers from pre-oriented lipid conformers selected to minimize steric overlaps while rejecting ring-piercing placements, and relaxes the assembled patch through soft-core followed by restrained minimization. Beyond the 41 built-in OpenMM Lipid21 templates, roughly 4,000 additional Lipid21 and lipid_ext species are generated on demand via tleap.

### Enhanced sampling and free energies

Molpylot provides a unified library of collective variables (spanning geometric and contact coordinates, conformational path variables^101^ and PyTorch-trained Deep-TICA coordinates^102^) evaluated identically in NumPy for trajectory analysis, as OpenMM custom forces during simulation and as PLUMED input files ^103^. Well-tempered metadynamics^42^ applies a gridbased Gaussian bias, writes PLUMED-compatible HILLS and COLVAR files, and reconstructs free-energy surfaces from summed hills or *c*(*t*) frame reweighting^104^ with block-averaged error estimates^105^. Constant-velocity steered MD is analyzed via Jarzynski’s equality^44^ and secondorder cumulant expansion; umbrella sampling uses WHAM^43^ with bootstrap uncertainties; and Gaussian-accelerated MD^106^ employs dual-boost potentials with cumulant reweighting^107^. RBFE campaigns are executed through OpenFE 1.13^108^ in a dedicated environment driven by Molpylot. Atom mappings are generated with Kartograf or Lomap^109,110^, perturbation graphs are built with OpenFE network planners, and each alchemical transformation is run using OpenFE’s hybrid-topology replica-exchange protocol (defaulting to OpenFF Sage 2.3.0/NAGL ligands, ff14SB, TIP3P, Gapsys soft-core potentials^111^, 11 λ windows and three independent repeats per leg). Relative free energies are estimated with MBAR^112^, audited for cycle closure, window overlap and replica-exchange mixing, and converted into maximum-likelihood ligand binding affinities using cinnabar^113^.

### Trajectory analysis, docking and ligand-based design

Forty-seven built-in analyses execute in memory on worker threads and attach their outputs to the parent entry. To characterize conformational ensembles, Molpylot combines Kabsch superposition^114^, essential-dynamics PCA^115^ and time-lagged independent component analysis (TICA)^116^ with kinetic and commute maps^117^ (the kinetic map validated against deeptime^118^), alongside Daura^119^ and *k*-means clustering, self-organizing maps^120,121^ and graph-neural-network embeddings^122^. Local structural and pocket properties are quantified via DSSP secondary-structure assignment^123^, Shrake–Rupley solvent-accessible surface area^124^, native-contact fractions^125^ and grid-based pocket cavity integration. Long-timescale kinetics and allosteric coupling are modeled through reversible Markov state models with PCCA+ coarse-graining^126^ and Chapman–Kolmogorov validation^127^, and through contact networks weighted by dynamic cross-correlation^128^ or linear mutual information^129^ to compute Brandes betweenness^130^ and optimal communication paths^131^. Membrane modules compute leaflet-resolved density profiles, bilayer thickness, water penetration, deuterium order parameters and drift-corrected lateral diffusion. Across all time series, equilibration times and statistical uncertainties are evaluated via autocorrelation analysis and Flyvbjerg–Petersen block averaging^34,36^. Interactive visualizations use PyQtGraph^132^, while static exports use Matplotlib^133^ with a color-blind-safe palette^134^.

For structureand ligand-based design, Molpylot integrates AutoDock Vina 1.2^135^ with Vina and Vinardo scoring^136^ and Meeko^137^ ligand preparation, supporting pH-dependent state enumeration, ensemble receptor docking, core-constrained posing, symmetry-aware RMSD evaluation and MM-GBSA rescoring in OBC2 implicit solvent^138^. Pharmacophores and protein–ligand interaction fingerprints can be derived from ligand series, complexes, apo pockets or trajectories, and ligand-based QSAR models (classical machine-learning regressors and Chemprop graph neural networks) are trained with scaffold-based cross-validation and applicability-domain estimation.

### Agent-driven preparation and analysis of the validation set

The scientist delegated the preparation and analysis of all 18 validation systems to Claude Code agents running Claude Opus 5.5 (medium effort) and supervised their work; the agents were restricted exclusively to the stand-alone molpylot-mcp server over stdio, with shell execution, file reading and writing, web access, sub-agents and all other MCP servers disabled in the client permissions (standing instructions, a representative prompt and the client configuration in Supplementary Note 2). During preparation, agents were instructed to discover tools via find_tools and search_docs, retain Molpylot’s default force fields and protocols unless a physical check or the system’s biology required an adjustment, resolve or explicitly justify every check warning, save a session project per system, and export self-contained job folders rather than launching simulations locally. Check findings, system-specific force-field settings and agent–scientist decisions are detailed in Supplementary Table 4, Extended Data Table 2 and Supplementary Sections 1–5.

Production MD jobs were executed on the CINECA Leonardo supercomputer as chained 24 h allocations on single NVIDIA A100 GPUs (8 CPU cores, 64 GB RAM), writing a checkpoint upon receiving SIGTERM 10 min before wall-time expiry and resuming automatically in the next job (achieving 422 ns day⁻¹ at a 2 fs time step for the 25,837-atom solvated ubiquitin system).

Trajectory and free-energy analyses were carried out by tool-confined agents operating on copies of the completed job folders. For each replica, the analysis agent used automated convergence detection on the backbone RMSD (plus ligand RMSD for complexes and bilayer area for membranes) to define the post-equilibration analysis window, and computed every property over both the equilibrated window and the full production trajectory (Supplementary Table 5 and Supplementary Data 3). Unless noted otherwise, reported uncertainties represent block-averaged standard errors within individual replicas and standard deviations of replica means across independent replicas.

Standard geometric criteria were applied across all systems: hydrogen bonds required a donor–acceptor distance ≤ 3.5 Å and a D–H⋯A angle ≥ 150° (≥ 120° in the analyses of PDB 1A28 and 4DFR and of the waters around the rhodopsin Schiff base; N⋯S ≤ 3.8 Å and ≥ 120° for the NH⋯S hydrogen bonds of rubredoxin); salt bridges required an O–N distance ≤ 4.0 Å; and water bridges required a water oxygen within 3.5 Å of polar atoms on both partners. Ligand heavy-atom RMSDs were calculated after aligning the backbone of residues within 5 Å of the reference ligand pose. In metalloproteins, a coordinating donor was classified as lost if its metal distance exceeded 3.0 Å in more than 10% of frames, and first-shell water coordination was evaluated within 2.8 Å. System-specific benchmarks used established literature definitions: the dihydrofolate reductase Met20 loop (residues 9–24) was classified against the occluded crystal structure and the closed reference 1RX2^139^ using a 2.0 Å backbone RMSD threshold; the

EGFR DFG-in conformation was assigned using Kincore distance criteria^140^; lipid bilayer areas, thicknesses and order parameters were compared against small-angle scattering and NMR measurements at matching temperatures^141–143^; and TYK2 RBFE campaigns were evaluated against experimental affinities^45^ via bootstrap RMSE, MUE, *R*² and Kendall τ.

### Validation of components

Individual computational engines and analysis modules were benchmarked against reference implementations (numerical comparisons in Extended Data Table 1). Across 25 random configurations per variable type, values of collective variables agree with PLUMED 2.9 to within 1.6 × 10⁻⁷ (relative) and with OpenMM custom forces to within 10⁻¹³. A 10 ns welltempered metadynamics run of alanine dipeptide in Molpylot reproduces the φ free-energy profile of PLUMED 2.9 driving the same OpenMM system within 0.17–0.25 kcal mol⁻¹ RMSD across two random seeds (smaller than the 0.31–0.33 kcal mol⁻¹ between-seed spread of either engine alone), and hill summation matches PLUMED sum_hills. Exported Amber and GROMACS topologies and coordinates reproduce single-point energy terms of sander to within 5 × 10⁻⁵ kJ mol⁻¹ and of GROMACS 2026 to within 8.1 × 10⁻⁴ kJ mol⁻¹; cofactor and heme topologies match tleap term by term.

In benchmarks of structure preparation and ligand chemistry across 216 diverse PDB entries, 212 structures completed automated preparation while four were halted as intended by pre-execution checks on incomplete dictionary ligands, and for 156 of the 158 free ligands the molecule in the prepared structure matched its PDB Chemical Component Dictionary definition in heavy atoms, bonds and bond orders (standard InChI without stereo and protonation layers). In redocking benchmarks across nine protein–ligand complexes (exhaustiveness 16, fixed seed), AutoDock Vina integration placed the top-ranked pose within 2.0 Å RMSD of the crystallographic pose for eight of nine targets. Finally, hydrogen-bond triplets and continuous autocorrelation lifetimes agree with MDAnalysis within 0.002, grid-based cavity volume integration requires 5.7 ms per frame on a laptop CPU for solvated ubiquitin, and OpenFE campaign aggregation reproduces openfe gather outputs.

### Use of a large language model in writing

A large language model was used to assist with drafting and editing the text of the manuscript and the Supplementary Information and with checking cross-references and citations. The authors directed this work, reviewed and revised all text, and take full responsibility for the content.

## Data availability

Structures were obtained from the Protein Data Bank under accession codes 1UBQ, 1A28, 4DFR, 1HSG, 3ERT, 1M17, 1ZAA, 3HS4, 1IRO, 1A6N, 4WB5, 6OIM, 1U19, 4EIY, 1RX2 and 1BL8. The TYK2 ligand series and its measured affinities are those of the published benchmark ^45^. The tool catalog, the physical-check rules, the per-replica results of the validation set, the lipid order parameters, and the per-ligand, per-edge and cycle-closure values of the TYK2 campaigns are provided as Supplementary Data 1–7. Other data supporting the findings of this study are available from the corresponding author upon reasonable request.

## Code availability

Molpylot is available at https://github.com/Molpylot-Lab/releases. Molpylot is free for noncommercial use under the PolyForm Strict License 1.0.0, with additional permissions to modify the software for internal use and, for anyone including commercial organizations, to install and run it in order to peer review a scientific work or to verify or reproduce published results. Commercial use requires a license from the University of Milano-Bicocca.

## Acknowledgments

We acknowledge ISCRA for awarding this project (IscrB_NR-DynAt) access to the LEONARDO supercomputer, owned by the EuroHPC Joint Undertaking, hosted by CINECA (Italy), on which the simulations of the validation set were carried out. This work received no specific funding.

## Author contributions

S.M.: conceptualization, methodology, software, validation, formal analysis, investigation, resources, data curation, visualization, writing – original draft, writing – review and editing, supervision and project administration. A.R.: software (testing, debugging and design suggestions), validation, investigation (molecular dynamics simulations) and writing – review and editing. L.C. and G.S.: software (testing and debugging), validation, investigation (molecular dynamics simulations) and writing – review and editing. All authors reviewed and approved the manuscript.

## Competing interests

Molpylot is free for noncommercial use; commercial use requires a license from the University of Milano-Bicocca, with which all authors are affiliated. The authors declare no other competing interests.

## Extended Data

**Extended Data Fig. 1.**
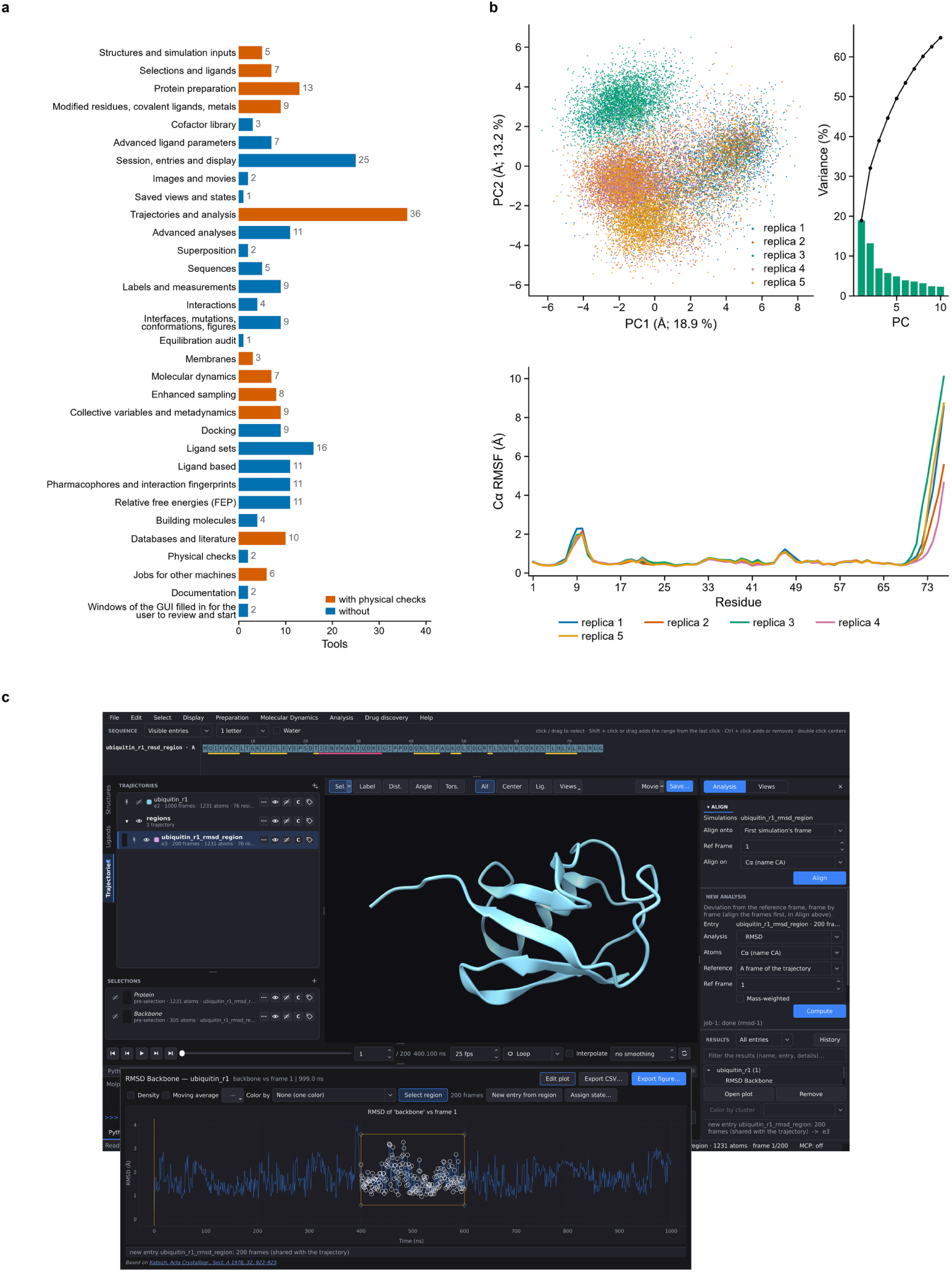
From a structure to a simulation, and a trajectory that can be navigated. **a**, The 32 tool areas of the Molpylot catalog in registration order from structures and preparation to jobs and documentation, with the number of tools in each area; the 11 areas holding tools wrapped by Molpylot automated physical checks are highlighted in orange. **b**, Native multi-replica analysis of ubiquitin (5 × 1 µs, 100–1000 ns): joint principal-component projection of Cα atoms (residues 1–70) colored by replica (PC1 18.9%, PC2 13.2% variance) and per-replica Cα RMSF profiles. **c**, Interactive linked plot-to-structure selection: brushing a region (400–600 ns) on the backbone RMSD plot of replica 1 creates a zero-copy 200-frame entry stored by frame index rather than duplicated coordinates, immediately listed in the session and displayed in the 3D viewport.

**Extended Data Fig. 2.**
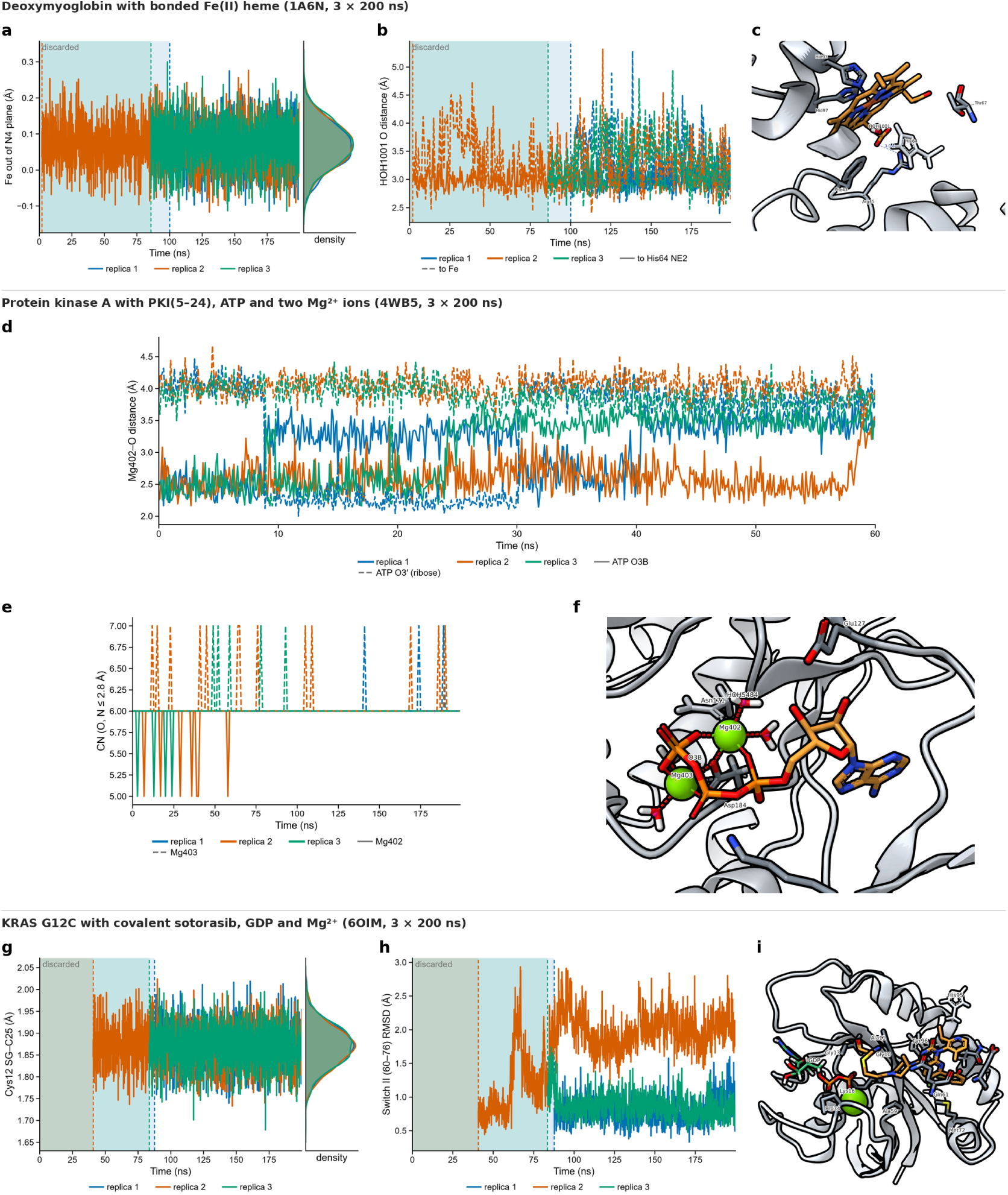
**Cofactor, phosphorylated and covalent validation complexes (1A6N, 4WB5 and 6OIM). a–c**, Deoxymyoglobin with the Amber bonded Fe(II) heme model and His64 set to HIE (1A6N, 3 × 200 ns): signed displacement of Fe from the porphyrin N₄ plane (positive toward proximal His93; crystal +0.36 Å) (**a**), distances from distal crystal water HOH1001 O to His64 Nε2 (solid) and Fe (dashed) (**b**), and representative 3D view of the heme pocket (replica 1, 199.6 ns) (**c**). **d–f**, Human protein kinase A catalytic subunit with PKI(5–24), ATP⁴⁻, three phosphorylated residues (pSer139, pThr197, pSer338) and two Li–Merz 12-6-4 Mg²⁺ ions (4WB5, 3 × 200 ns): Mg402–O3B (solid) and Mg402–ribose O3′ (dashed) distances over 0–60 ns showing permanent O3B departure and bulk-water substitution in all three replicas (**d**), Mg402 (solid) and Mg403 (dashed) coordination numbers (O, N ≤ 2.8 Å) (**e**), and 3D view of the active site (replica 1, 199.1 ns) showing replacing water HOH5484 on Mg402 (**f**). **g–i**, KRAS G12C covalently bound to sotorasib with GDP³⁻ and 12-6-4 Mg²⁺ (6OIM, 3 × 200 ns): Cys12 Sγ–sotorasib C25 covalent bond length (**g**), Switch II (residues 60–76) backbone RMSD fitted on the protein core outside the switches showing the replica 2 conformational shift at 62–66 ns (**h**), and 3D view of the switch-II pocket and nucleotide site (replica 1, 199.6 ns) (**i**). Shaded intervals mark per-replica discarded equilibration windows.

**Extended Data Fig. 3.**
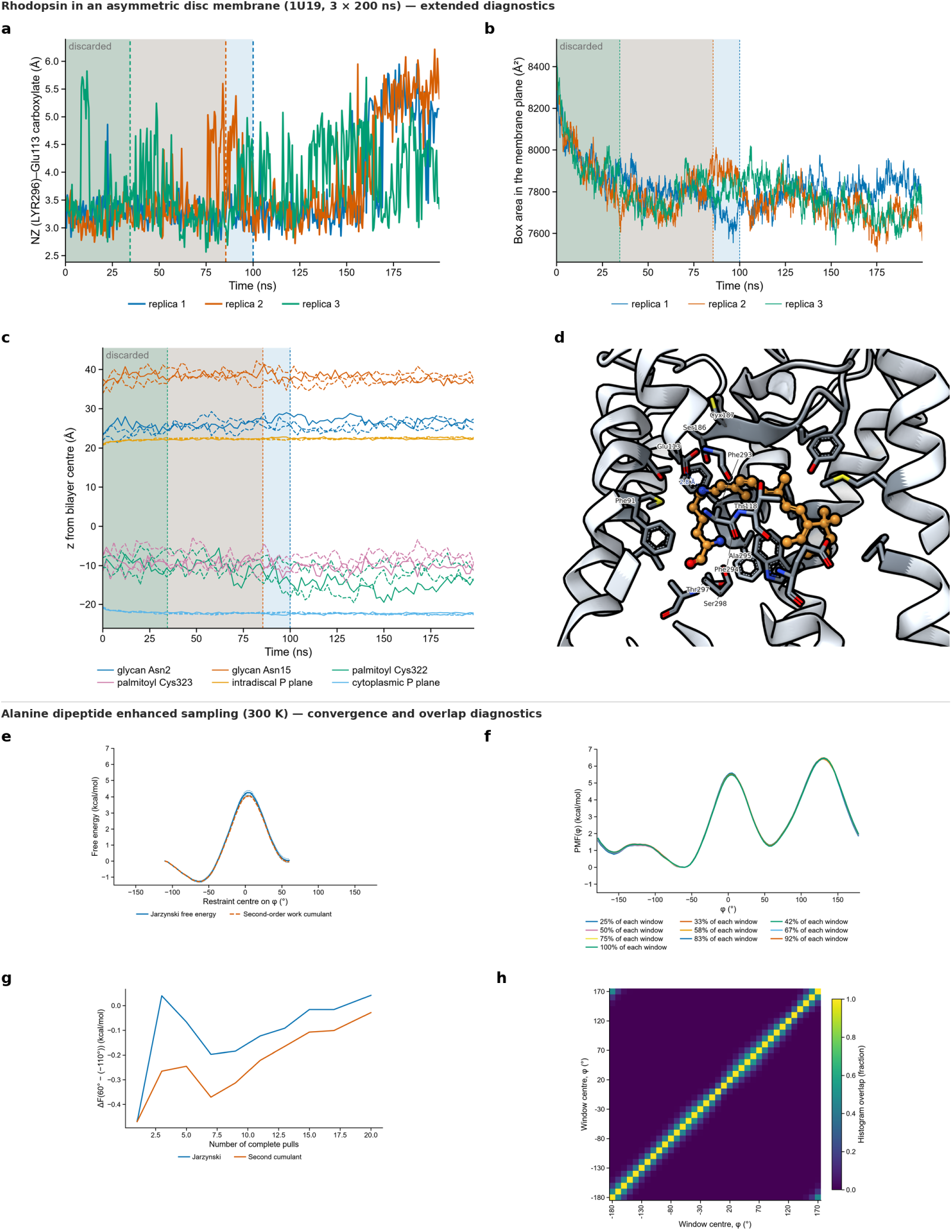
**Extended diagnostics for asymmetric rhodopsin (1U19) and alanine dipeptide enhanced sampling. a–d**, Bovine rhodopsin in an agent-configured asymmetric disc membrane (1U19, 3 × 200 ns; complementing Fig. 3a–c): Schiff-base Nζ(LYR296)–Glu113 carboxylate distance showing intermittent opening compensated by crystal-water bridges (**a**), membrane-plane box area over 0–200 ns showing the initial 6–7% contraction and slow area fluctuations (**b**), *z*-coordinates relative to the bilayer midplane for both intradiscal *N*-glycans, both cytoplasmic palmitoyl chains and the two phosphate planes (replica 1 solid, replicas 2–3 dashed) (**c**), and 3D view of the 11-*cis* retinal pocket (replica 3, 199.6 ns) (**d**). **e–h**, Alanine dipeptide in OPC water at 300 K (complementing Fig. 4): steered-MD Jarzynski free-energy profile versus second-order cumulant expansion across −110° to +60° (**e**), umbrella-sampling PMF stability across growing fractions (25–100%) of each 3.75 ns post-transient window (**f**), steered-MD Δ*F* (60° − (−110°)) convergence over 20 pulls (**g**), and neighboring-window histogram overlap matrix across all 36 umbrella windows after discarding the initial 1.25 ns transient (all adjacent overlaps 0.27–0.52) (**h**).

**Extended Data Fig. 4.**
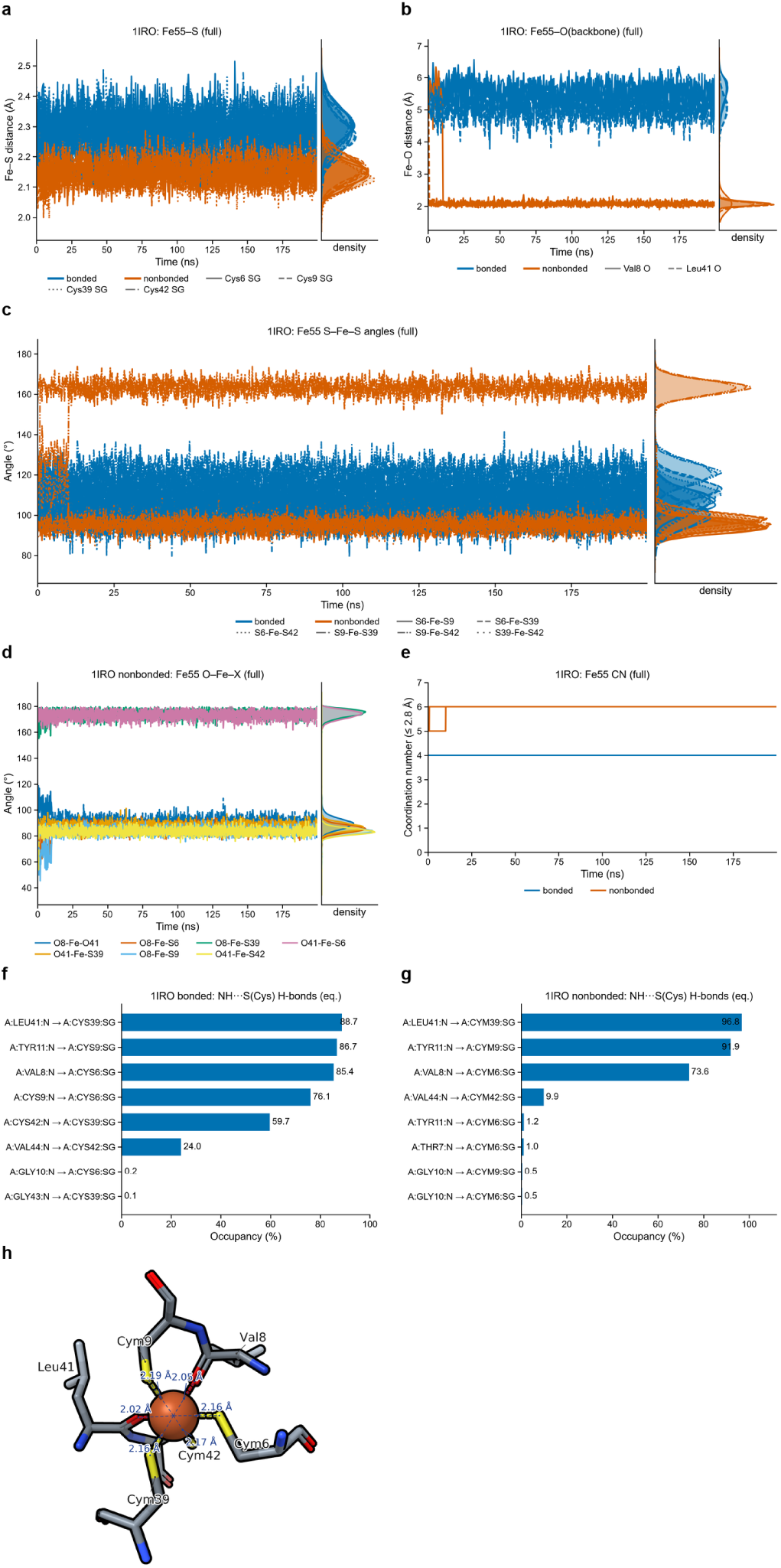
Bonded versus non-bonded Fe(III) site in oxidized rubredoxin (1IRO). Bonded high-spin Fe(III) site (Seminario B3LYP/6-31G* force constants and RESP charges; blue) versus non-bonded Li– Merz 12-6 Fe³⁺ control (orange) over 3 × 200 ns per model. **a**, Fe55–Cys Sγ distances across the whole production trajectories; the bonded model maintains all four Fe–S bonds within 0.06 Å of the 1.1 Å crystal structure, whereas the non-bonded ion compresses Fe–S by ∼0.1 Å. **b**, Distances from Fe55 to the backbone carbonyl oxygens of Val8 and Leu41, which stay 5.1–5.5 Å away on average (s.d. about 0.4 Å) in the bonded model but coordinate the non-bonded iron at 2.07–2.08 Å. **c**, S–Fe–S angles across the full trajectories; the bonded site stays tetrahedral, whereas S9–Fe–S42 opens to 163° in the non-bonded control. **d**, *Trans* O–Fe–S angles (173–174°) in the collapsed octahedral Fe(S)₄(O)₂ non-bonded site. **e**, Iron coordination number (O, N, S ≤ 2.8 Å): constant at 4.00 in the bonded model versus 6.00 in the non-bonded control. **f**,**g**, Protective backbone NH···Sγ hydrogen-bond occupancies in the post-equilibration windows for the bonded (**f**) and non-bonded (**g**) models; Cys9→S6 and Cys42→S39 are completely lost in the non-bonded control. **h**, Representative 3D structure of the non-bonded iron site (replica 2, 150.0 ns) showing octahedral collapse onto Val8 O and Leu41 O.

---

**Extended Data Table 1.**
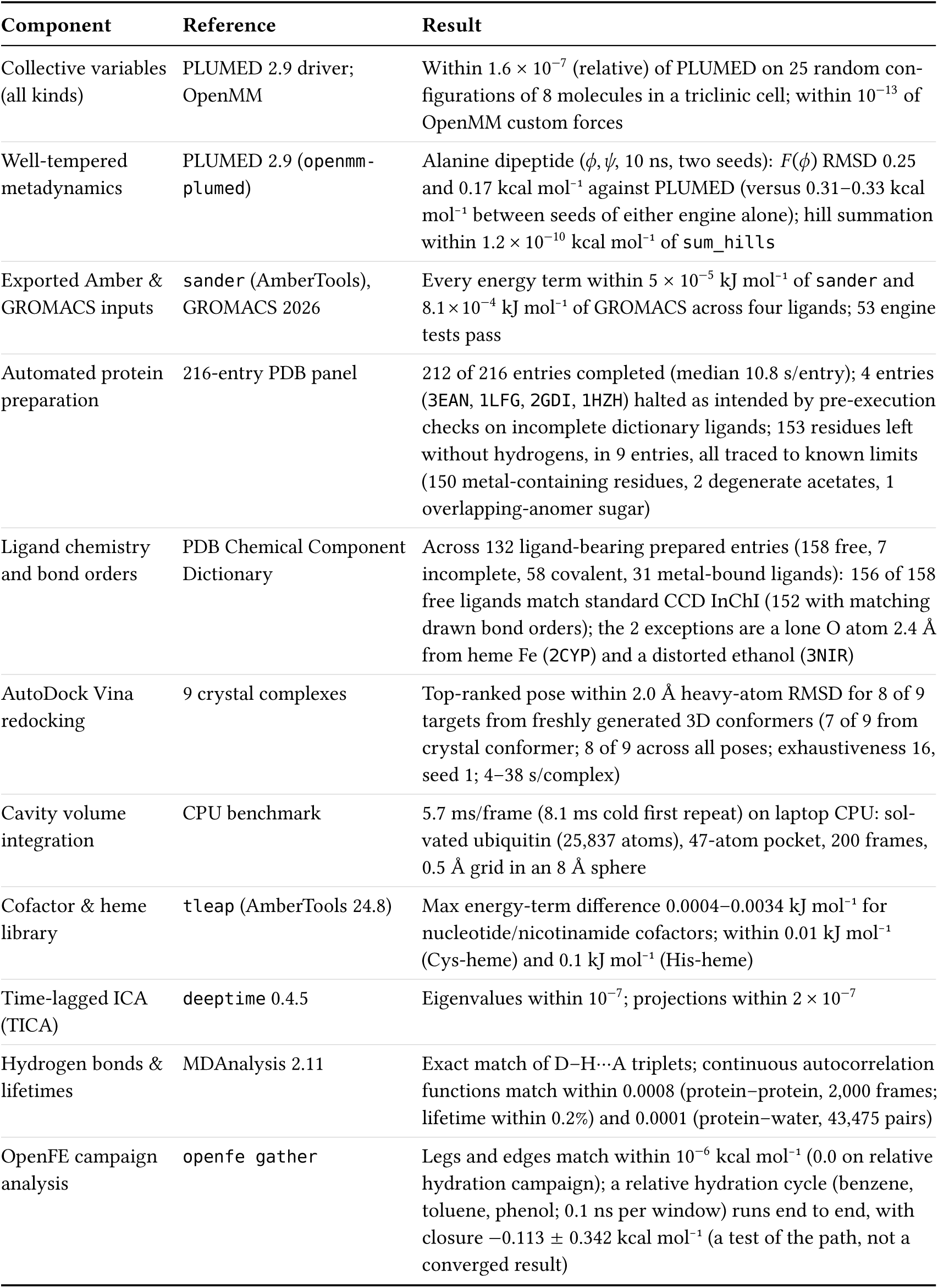
Validation of Molpylot components against reference implementations and experimental benchmarks.

| **Component** | **Reference** | **Result** |
|---|---|---|
| Collective variables (all kinds) | PLUMED 2.9 driver; OpenMM | Within 1.6 × 10<sup>−7</sup> (relative) of PLUMED on 25 random con-figurations of 8 molecules in a triclinic cell; within 10<sup>−13</sup> of OpenMM custom forces |
| Well-tempered metadynamics | PLUMED 2.9 (openmm-plumed) | Alanine dipeptide (*ϕ*, *ψ*, 10 ns, two seeds): *F*(*ϕ*) RMSD 0.25 and 0.17 kcal mol<sup>−1</sup> against PLUMED (versus 0.31–0.33 kcal mol<sup>−1</sup> between seeds of either engine alone); hill summation within 1.2 × 10<sup>−10</sup> kcal mol<sup>−1</sup> of sum_hills |
| Exported Amber & GROMACS inputs | sander (AmberTools), GROMACS 2026 | Every energy term within 5 × 10<sup>−5</sup> kJ mol<sup>−1</sup> of sander and 8.1 × 10<sup>−4</sup> kJ mol<sup>−1</sup> of GROMACS across four ligands; 53 engine tests pass |
| Automated protein preparation | 216-entry PDB panel | 212 of 216 entries completed (median 10.8 s/entry); 4 entries (3EAN, 1LFG, 2GDI, 1HZH) halted as intended by pre-execution checks on incomplete dictionary ligands; 153 residues left without hydrogens, in 9 entries, all traced to known limits (150 metal-containing residues, 2 degenerate acetates, 1 overlapping-anomer sugar) |
| Ligand chemistry and bond orders | PDB Chemical Component Dictionary | Across 132 ligand-bearing prepared entries (158 free, 7 incomplete, 58 covalent, 31 metal-bound ligands): 156 of 158 free ligands match standard CCD InChI (152 with matching drawn bond orders); the 2 exceptions are a lone O atom 2.4 Å from heme Fe (2CYP) and a distorted ethanol (3NIR) |
| AutoDock Vina redocking | 9 crystal complexes | Top-ranked pose within 2.0 Å heavy-atom RMSD for 8 of 9 targets from freshly generated 3D conformers (7 of 9 from crystal conformer; 8 of 9 across all poses; exhaustiveness 16, seed 1; 4–38 s/complex) |
| Cavity volume integration | CPU benchmark | 5.7 ms/frame (8.1 ms cold first repeat) on laptop CPU: sol-vated ubiquitin (25,837 atoms), 47-atom pocket, 200 frames, 0.5 Å grid in an 8 Å sphere |
| Cofactor & heme library | tleap (AmberTools 24.8) | Max energy-term difference 0.0004–0.0034 kJ mol<sup>−1</sup> for nucleotide/nicotinamide cofactors; within 0.01 kJ mol<sup>−1</sup> (Cys-heme) and 0.1 kJ mol<sup>−1</sup> (His-heme) |
| Time-lagged ICA (TICA) | deeptime 0.4.5 | Eigenvalues within 10<sup>−7</sup>; projections within 2 × 10<sup>−7</sup> |
| Hydrogen bonds & lifetimes | MDAnalysis 2.11 | Exact match of D–H···A triplets; continuous autocorrelation functions match within 0.0008 (protein–protein, 2,000 frames; lifetime within 0.2%) and 0.0001 (protein–water, 43,475 pairs) |
| OpenFE campaign analysis | openfe gather | Legs and edges match within 10<sup>−6</sup> kcal mol<sup>−1</sup> (0.0 on relative hydration campaign); a relative hydration cycle (benzene, toluene, phenol; 0.1 ns per window) runs end to end, with closure −0.113 ± 0.342 kcal mol<sup>−1</sup> (a test of the path, not a converged result) |

**Extended Data Table 2.**
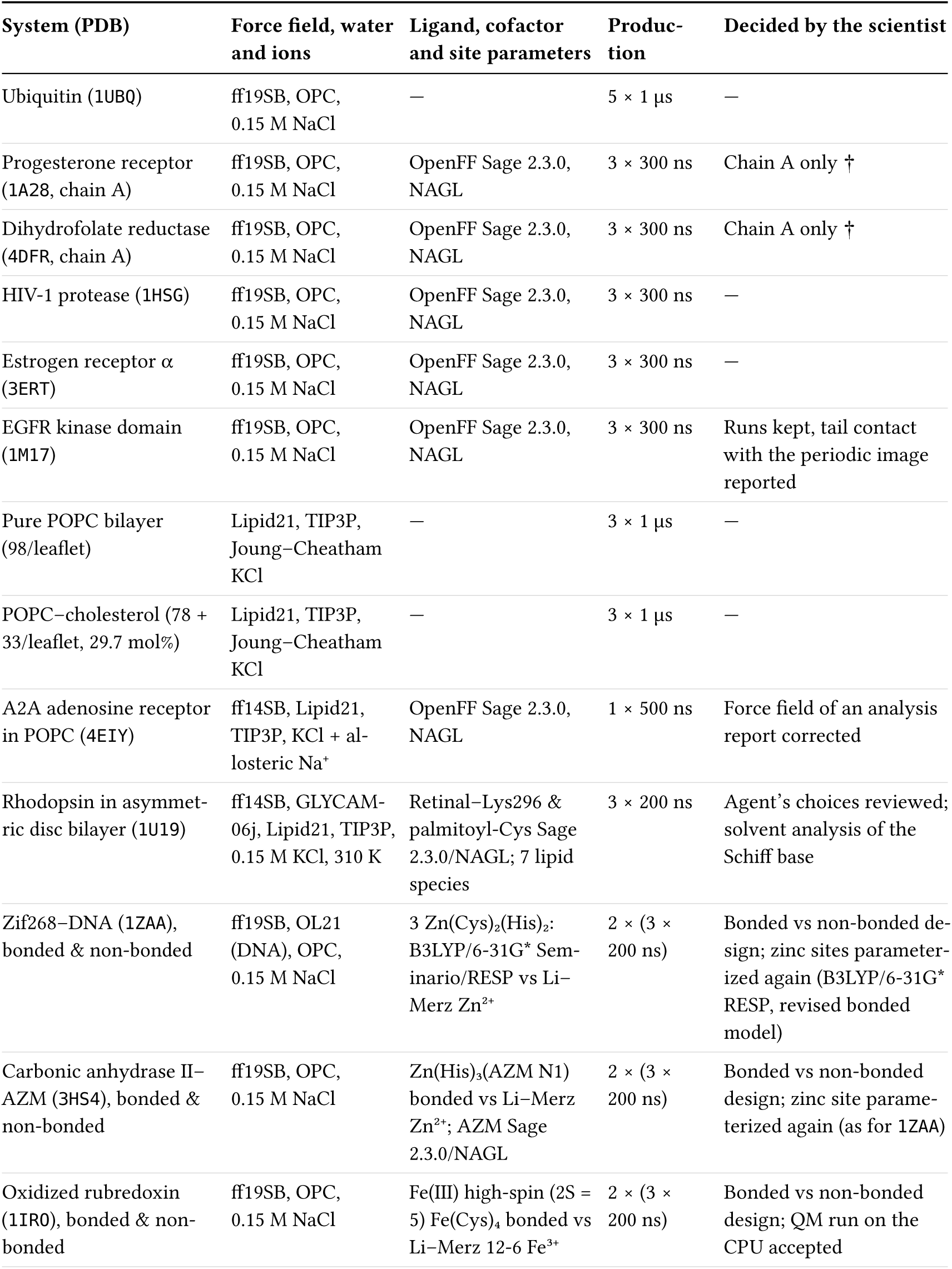
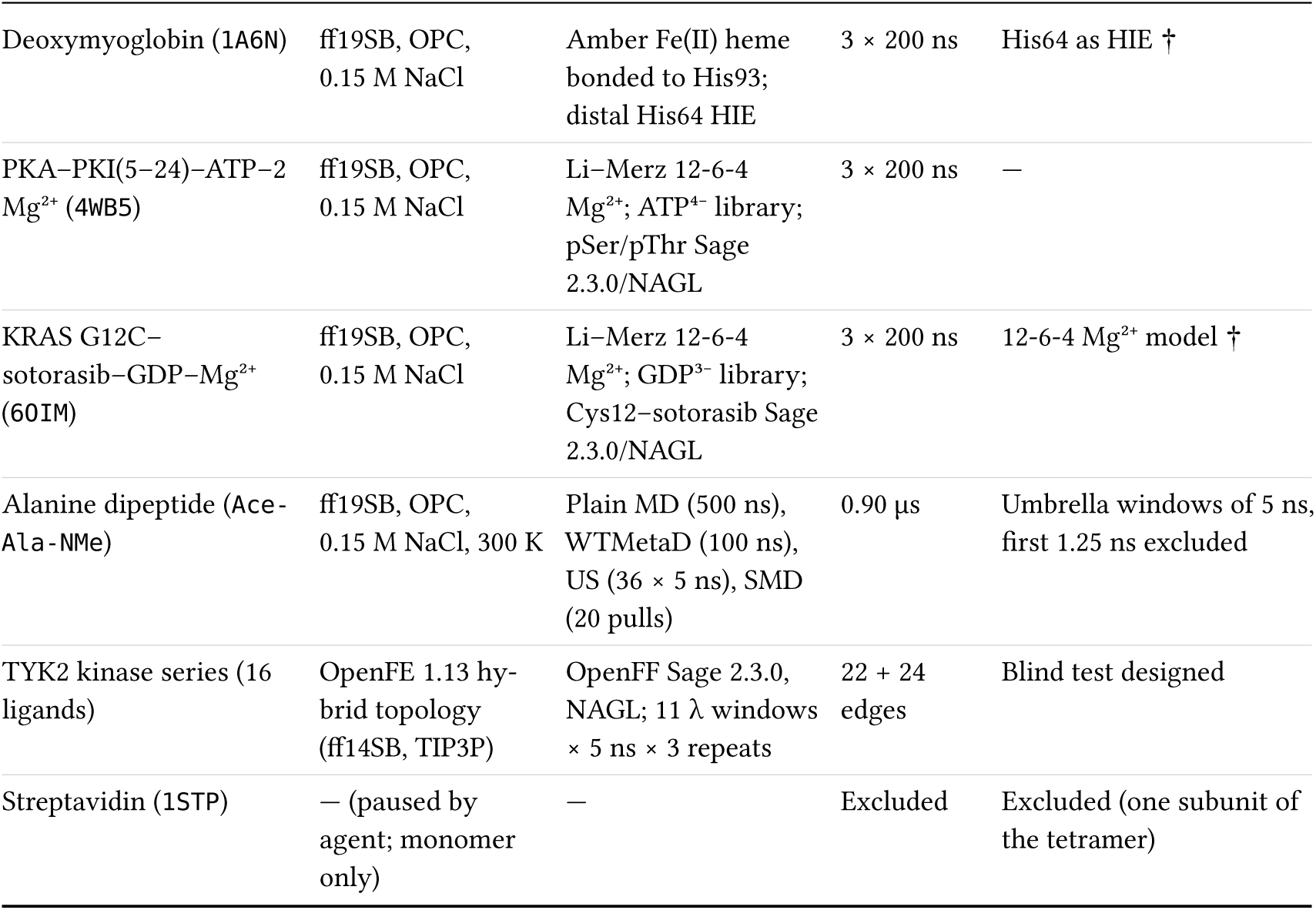
Overview of the 18-system agent-prepared validation suite. The scientist delegated the preparation and analysis of all systems to Claude Code agents running Claude Opus 5.5 (medium effort), restricted to Molpylot’s MCP tools, and supervised their work; the last column lists the decisions the scientist took on the study design, during preparation and during analysis (details in Supplementary Table 4). Those marked † answered a question an agent raised during preparation; the other 14 systems needed no scientific input at that stage. Total production time: 22.0 µs across the 57 biomolecular MD trajectories, plus 0.90 µs of alanine dipeptide across four sampling methods and 276 alchemical leg-repeats across two TYK2 RBFE campaigns.

| System (PDB) | Force field, water and ions | Ligand, cofactor and site parameters | Produc-tion | Decided by the scientist |
|---|---|---|---|---|
| Ubiquitin (1UBQ) | ff19SB, OPC, 0.15 M NaCl | — | 5 × 1 µs | — |
| Progesterone receptor (1A28, chain A) | ff19SB, OPC, 0.15 M NaCl | OpenFF Sage 2.3.0, NAGL | 3 × 300 ns | Chain A only † |
| Dihydrofolate reductase (4DFR, chain A) | ff19SB, OPC, 0.15 M NaCl | OpenFF Sage 2.3.0, NAGL | 3 × 300 ns | Chain A only † |
| HIV-1 protease (1HSG) | ff19SB, OPC, 0.15 M NaCl | OpenFF Sage 2.3.0, NAGL | 3 × 300 ns | — |
| Estrogen receptor α (3ERT) | ff19SB, OPC, 0.15 M NaCl | OpenFF Sage 2.3.0, NAGL | 3 × 300 ns | — |
| EGFR kinase domain (1M17) | ff19SB, OPC, 0.15 M NaCl | OpenFF Sage 2.3.0, NAGL | 3 × 300 ns | Runs kept, tail contact with the periodic image reported |
| Pure POPC bilayer (98/leaflet) | Lipid21, TIP3P, Joung–Cheatham KCl | — | 3 × 1 µs | — |
| POPC–cholesterol (78 + 33/leaflet, 29.7 mol%) | Lipid21, TIP3P, Joung–Cheatham KCl | — | 3 × 1 µs | — |
| A2A adenosine receptor in POPC (4EIY) | ff14SB, Lipid21, TIP3P, KCl + al-losteric Na<sup>+</sup> | OpenFF Sage 2.3.0, NAGL | 1 × 500 ns | Force field of an analysis report corrected |
| Rhodopsin in asymmet-ric disc bilayer (1U19) | ff14SB, GLYCAM-06j, Lipid21, TIP3P, 0.15 M KCl, 310 K | Retinal–Lys296 & palmitoyl-Cys Sage 2.3.0/NAGL; 7 lipid species | 3 × 200 ns | Agent’s choices reviewed; solvent analysis of the Schiff base |
| Zif268–DNA (1ZAA), bonded & non-bonded | ff19SB, OL21 (DNA), OPC, 0.15 M NaCl | 3 Zn(Cys)<sub>2</sub>(His)<sub>2</sub>: B3LYP/6-31G<sup>*</sup> Sem-inario/RESP vs Li–Merz Zn<sup>2+</sup> | 2 × (3 × 200 ns) | Bonded vs non-bonded de-sign; zinc sites parameter-ized again (B3LYP/6-31G<sup>*</sup> RESP, revised bonded model) |
| Carbonic anhydrase II–AZM (3HS4), bonded & non-bonded | ff19SB, OPC, 0.15 M NaCl | Zn(His)<sub>3</sub>(AZM N1) bonded vs Li–Merz Zn<sup>2+</sup>; AZM Sage 2.3.0/NAGL | 2 × (3 × 200 ns) | Bonded vs non-bonded design; zinc site parame-terized again (as for 1ZAA) |
| Oxidized rubredoxin (1IRO), bonded & non-bonded | ff19SB, OPC, 0.15 M NaCl | Fe(III) high-spin (2S = 5) Fe(Cys)<sub>4</sub> bonded vs Li–Merz 12-6 Fe<sup>3+</sup> | 2 × (3 × 200 ns) | Bonded vs non-bonded design; QM run on the CPU accepted |
| Deoxymyoglobin (1A6N) | ff19SB, OPC, 0.15 M NaCl | Amber Fe(II) heme bonded to His93; distal His64 HIE | 3 × 200 ns | His64 as HIE † |
| PKA–PKI(5–24)–ATP–2 Mg<sup>2+</sup> (4WB5) | ff19SB, OPC, 0.15 M NaCl | Li–Merz 12-6-4 Mg<sup>2+</sup>; ATP<sup>4−</sup> library; pSer/pThr Sage 2.3.0/NAGL | 3 × 200 ns | — |
| KRAS G12C–sotorasib–GDP–Mg<sup>2+</sup> (6OIM) | ff19SB, OPC, 0.15 M NaCl | Li–Merz 12-6-4 Mg<sup>2+</sup>; GDP<sup>3−</sup> library; Cys12–sotorasib Sage 2.3.0/NAGL | 3 × 200 ns | 12-6-4 Mg<sup>2+</sup> model † |
| Alanine dipeptide (Ace-Ala-NMe) | ff19SB, OPC, 0.15 M NaCl, 300 K | Plain MD (500 ns), WTMetaD (100 ns), US (36 × 5 ns), SMD (20 pulls) | 0.90 µs | Umbrella windows of 5 ns, first 1.25 ns excluded |
| TYK2 kinase series (16 ligands) | OpenFE 1.13 hy-brid topology (ff14SB, TIP3P) | OpenFF Sage 2.3.0, NAGL; 11 λ windows × 5 ns × 3 repeats | 22 + 24 edges | Blind test designed |
| Streptavidin (1STP) | — (paused by agent; monomer only) | — | Excluded | Excluded (one subunit of the tetramer) |

## Notes

https://github.com/Molpylot-Lab/releases

