## Supplementary Material for "Molpylot: a shared workspace for human–AI collaboration in molecular simulation"

#### Guide to Supplementary Material, Extended Data and Supplementary Data

This document contains summary tables, complementary diagnostic figures, system-level notes and methodological details supporting the main manuscript. Large machine-readable catalogs and per-replica numerical tables are provided as separate **Supplementary Data** CSV files.

| Resource | Content |
|---|---|
| **Extended Data Figs. 1–4** (with the main article) | Platform overview and linked brushing (ED Fig. 1); heme, phosphorylated Mg-ATP kinase and covalent KRAS G12C complexes (ED Fig. 2); non-redundant diagnostics for rhodopsin and alanine dipeptide (ED Fig. 3); rubredoxin Fe(III) bonded vs non-bonded comparison (ED Fig. 4). |
| **Extended Data Tables 1–2** (with the main article) | Component validation benchmarks against reference engines (ED Table 1); 18-system validation suite overview (ED Table 2). |
| **Supplementary Tables 1–7** | Tool catalog summary by area; exposed physical-check summary by family; MCP profile context sizes; check findings and the scientist's decisions in the validation campaign; master quantitative summary of all 18 systems; quantum-chemical models of the bonded metal sites; TYK2 RBFE campaign summary. |
| **Supplementary Figs. 1–3** | Composite diagnostics for soluble/protein–ligand complexes (SFig. 1), lipid bilayers and A2A receptor (SFig. 2), and extended zinc-site diagnostics for 1ZAA and 3HS4 (SFig. 3). |
| **Supplementary Sections 1–5** | Concise setup exceptions, observables and agent–scientist interactions across the 18 validation systems. |
| **Supplementary Videos 1–3** | Building a membrane-protein system through natural language in a live Molpylot session (KcsA, PDB 1BL8; Video 1); interactive plots linked to the 3D structure (Video 2); conformational landscapes, frame selection and coordinated residues (Video 3). |
| **Supplementary Notes 1–2** | Security model for MCP tool execution (Note 1); tool-confined agent configuration, standing rules and representative biological brief (Note 2). |
| **Supplementary Data 1–7** (`.csv`) | Complete 260-tool catalog (SD1); 80 exposed physical-check codes (SD2); analyzed-window and whole-production values of every MD and enhanced-sampling system, per replica where the analysis is per replica (SD3); per-carbon lipid $-S_{CD}$ profiles (SD4); TYK2 per-ligand $\Delta G$ (SD5), per-edge $\Delta\Delta G$ (SD6) and cycle closures (SD7). |

**Supplementary Table 1 | Molpylot tool catalog by functional area.** Summary of the 32 functional areas registered by the Molpylot MCP server. Of the 260 exposed tools, 256 belong to the unified catalog shared identically by the GUI, built-in Python console and MCP server, and 4 (`find_tools`, `get_job`, `list_jobs`, `cancel_job`) provide MCP-specific tool discovery and background-job tracking. The last column marks the 11 areas holding the 22 tools wrapped by physical checks (in area 28, `fetch_alphafold_model`); the two tools of area 29 run and configure the checks on demand. The complete per-tool catalog with descriptions is provided in `Supplementary_Data_1_tool_catalog.csv`.

| # | Functional area | Tools | Tools wrapped by physical checks |
|---|---|---|---|
| 1 | Structures and simulation inputs | 5 | Yes |
| 2 | Selections and ligands | 7 | Yes |
| 3 | Protein preparation | 13 | Yes |
| 4 | Modified residues, covalent ligands, metals | 9 | Yes |
| 5 | Cofactor library | 3 | — |
| 6 | Advanced ligand parameters | 7 | — |
| 7 | Session, entries and display | 25 | — |
| 8 | Images and movies | 2 | — |
| 9 | Saved views and states | 1 | — |
| 10 | Trajectories and analysis | 36 | Yes |
| 11 | Advanced analyses: correlation networks, distance fluctuations, convergence, pores, Markov models, lipids | 11 | — |
| 12 | Superposition | 2 | — |
| 13 | Sequences | 5 | — |
| 14 | Labels and measurements | 9 | — |
| 15 | Interactions | 4 | — |
| 16 | Interfaces, mutations, conformations, figures | 9 | — |
| 17 | Equilibration audit | 1 | — |
| 18 | Membranes | 3 | Yes |
| 19 | Molecular dynamics | 7 | Yes |
| 20 | Enhanced sampling | 8 | Yes |
| 21 | Collective variables and metadynamics | 9 | Yes |
| 22 | Drug discovery: docking (AutoDock Vina) | 9 | — |
| 23 | Ligand sets: ligands, their poses and the calculations behind them | 16 | — |
| 24 | Drug discovery: ligand based (QSAR, screening) | 11 | — |
| 25 | Drug discovery: pharmacophores and interaction fingerprints | 11 | — |
| 26 | Relative free energies (FEP) | 11 | — |
| 27 | Building molecules | 4 | — |
| 28 | Databases and literature | 10 | Yes |
| 29 | Physical checks (`check_system`, `configure_checks`) | 2 | — |
| 30 | Jobs for other machines (`write_job`, `inspect_job`, ...) | 6 | Yes |

*(continued)*

| # | Functional area | Tools | Tools wrapped by physical checks |
|---|---|---|---|
| 31 | Documentation (`search_docs`, `find_tools`) | 2 | — |
| 32 | Windows of the GUI filled in for the user to review and start (`get_module_defaults`, `prefill_module`) | 2 | — |
| | **Total exposed over MCP** | **260** | **22 wrapped tools in 11 areas** |

**Supplementary Table 2 | Summary of exposed physical-check rules by family.** Overview of the 80 exposed physical-check finding codes across 12 families, wrapping 22 tools (11 with pre-execution input rules), on-demand `check_system`, and post-run `inspect_job(recheck=True)`. Five codes of the code base are not listed: four apply only to modules not yet exposed, and `open_shell`, which the exposed tools never emit. Four codes of finished runs are composed at run time (`equilibration_drifting`, `equilibration_unstable`, `production_drifting`, `production_unstable`) and are listed. Complete code-by-code messages, tool bindings and remediation text are provided in `Supplementary_Data_2_physical_checks.csv`.

| Check family | Codes (E / W / I) | Timing | Key physical criteria and representative findings |
|---|---|---|---|
| Structure: coordinates & geometry | 12 (5 / 5 / 2) | Pre / post | Overlapping atoms (< 0.8 Å heavy, < 0.5 Å any; error), non-finite coordinates (error), diverged minimization (error), broken bonds (< 1.1 or > 1.9 Å), severe clashes (0.8–1.6 Å), D-amino acids, water clashes (< 2.2 Å), non-proline *cis* peptides, hydrogens that could not be added during preparation (error) |
| Structure: chains & components | 12 (3 / 8 / 1) | Pre / post | Incomplete ligands or modified residues vs PDB CCD (error; blocks until scientist decides), missing heavy atoms, chain breaks, stretched peptide bonds, inverted stereocenters, disulfide erroneously bridging metal cysteines, unsupported modified nucleotides, a hydrogen on a metal-binding atom (< 2.0 Å from the metal; error) |
| Structure: parameters | 4 (0 / 4 / 0) | Pre | Unparameterized ligands or modified residues, crystallization additives retained, metal sites left as non-bonded ions |
| Periodic box | 5 (2 / 3 / 0) | Pre / post | Degenerate box or width < 2× cutoff (error), width within 6% of 2× cutoff, non-neutral net charge under PME, solute approaching its own periodic image |
| Simulation inputs | 8 (1 / 7 / 0) | Pre / post | Pulling/umbrella target distance exceeding half box width (error), small padding (< 8 Å), unusual temperature (< 250 or > 400 K) or pressure (> 10 bar), time step of 2–2.5 fs without HMR, NVT from unrelaxed density, funnel CV mismatch |
| Finished runs | 13 (1 / 9 / 3) | Post / recheck | Non-finite potential energy (error), mean temperature off thermostat target, unusual solvent density, periodic-image contact during trajectory, large Cα RMSD drift, equilibration or production log drift/instability |
| Metal sites: plans & parameters | 4 (0 / 4 / 0) | Pre | Planned but unparameterized site, stale parameters after plan change, failed QM/geometry validation, unbonded metal alongside bonded sites |

*(continued)*

| Check family | Codes (E / W / I) | Timing | Key physical criteria and representative findings |
|---|---|---|---|
| Metal sites along a run | 4 (0 / 2 / 2) | Post / recheck | Coordinating donor leaving cutoff (> 5% of frames), non-donor atom entering first shell (> 5% of frames) without bonded terms, intact coordination summary |
| Membranes | 3 (0 / 2 / 1) | Post / recheck | Bilayer area converged (reports per-replica equilibration start), area still drifting, or convergence not established (warns against starting enhanced sampling) |
| Enhanced sampling & free energies | 7 (0 / 6 / 1) | Post / recheck | Umbrella window overlap gaps, unsampled CV intervals, off-center windows, few SMD pulls or wide work spread ($\sigma \gg k_B T$), wide/anharmonic GaMD boost distribution |
| Ligand parameters | 6 (2 / 4 / 0) | Post | Partial-charge sum mismatch (error), impossible electron/spin multiplicity (error), extreme partial charges ($\lvert q \rvert > 1.3\,e$), poor RESP ESP fit, ligand lacking hydrogens or bound to unbonded metal |
| Trajectory analysis | 2 (0 / 2 / 0) | Post | Frame-to-frame RMSD jumps (unimaged periodic boundary crossings), extreme RMSD drift |
| **Total exposed** | **80 (14 / 56 / 10)** | | Under default `strict` policy, error-level input findings block execution and job export |

**Supplementary Table 3 | MCP tool profiles and model context footprint.** Compact JSON sizes of `tools/list` per built-in profile. “Read by model” counts tool names, titles, descriptions and input schemas. Server instructions occupy 3,831 characters (live embedded GUI session) or 3,972 characters (stand-alone headless server) against a 4,000-character cap. Seven core tools (`find_tools`, `search_docs`, `list_entries`, `get_system_summary`, `get_job`, `list_jobs`, `cancel_job`) are present in every profile.

| Profile | Tools | `tools/list` (chars) | Read by model (chars) | Output schemas (chars) |
|---|---|---|---|---|
| `full` (all tools) | 260 | 609,997 | 559,586 | 21,638 |
| `essential` (structures, prep, display, selections, measurements) | 80 | 131,399 | 116,181 | 6,517 |
| `simulation` (`essential` + ligand/metal params, membranes, MD, enhanced sampling) | 127 | 288,743 | 264,317 | 10,530 |
| `analysis` (`essential` + trajectory & advanced analyses, interfaces, sequences) | 142 | 285,464 | 258,184 | 11,664 |
| `drug_design` (`essential` + docking, ligand sets, QSAR, pharmacophores, FEP, DBs) | 152 | 302,185 | 272,838 | 12,564 |

**Supplementary Table 4 | Physical-check findings and the scientist's decisions during validation.** Distinct `(code, message)` pre-execution findings encountered during the preparation of the validation systems. Rows marked (*) come from the first parameterization of the zinc sites, before the scientist had them parameterized again, with B3LYP/6-31G* RESP charges and a revised bonded model.

| System | Finding code | Level | Reported condition | Outcome |
|---|---|---|---|---|
| 4DFR | `broken_bonds` | Warning | 3 C–O bonds at 1.08–1.09 Å | Resolved by preparation minimization |
| 4DFR | `incomplete_residues` | Warning | 9 residues missing heavy atoms | Resolved (rebuilt) |
| 4DFR | `cis_peptides` (×3) | Info | Gly95–Gly96 *cis* peptide in deposited copies | Kept (real crystallographic *cis* peptide) |
| 1M17 | `chain_breaks` | Warning | His964 ⟶ Leu977 break (4.6 Å) | Resolved (12-residue loop 965–976 built) |
| 3ERT | `incomplete_residues` | Warning | 12 incomplete residues | Resolved (rebuilt; backbone-less Pro552 removed) |
| 1A28 | `incomplete_residues` | Warning | 5 incomplete residues | Resolved (rebuilt) |
| 4EIY | `chain_breaks` | Warning | Lys1042 ⟶ Met1058 break (12.4 Å) | Resolved (BRIL loop 1043–1057 built) |
| 4EIY | `incomplete_residues` | Warning | 7 incomplete residues | Resolved (rebuilt) |
| 4EIY | `ligands_without_parameters` | Warning | ZMA unparameterized | Resolved (parameterized with Sage 2.3.0/NAGL) |
| 4EIY | `solute_sees_its_image` | Warning | 3.2 Å from periodic image in crystal cell | Resolved (replaced by membrane box) |
| 3HS4 | `severe_clashes`* / `water_clashes` | Warning | Clashing crystal waters (1.52–2.03 Å) | Resolved (removed by water filter, 13–22 Å from Zn) |
| 3HS4 | `charges_large`* | Warning | Acetazolamide sulfonyl S1 at +1.55 *e* | Accepted with reason (expected for sulfonyl S) |
| 3HS4 | `ligand_bound_to_metal`* | Warning | AZM bound to unbonded metal | Resolved in bonded model; accepted in non-bonded control |
| 1ZAA, 3HS4, 1IRO | `metal_sites_nonbonded` (×3) | Warning | Non-bonded $Zn^{2+}$ / $Fe^{3+}$ models | Accepted with reason (non-bonded control runs) |
| 1IRO | `water_clashes` | Warning | 2 waters at 2.02–2.11 Å | Resolved (removed by default preparation) |
| Metal jobs (×2) | `constant_volume_density_low` | Warning | Starting-box density 0.950–0.963 g $mL^{-1}$ | Accepted (applies only to initial 15 ps NVT before NPT) |
| 1A6N | `water_clashes` | Warning | HOH1130 2.14 Å from Glu41 Oε1 | Resolved (10.8 Å from heme; removed) |
| 4WB5 | `water_clashes` | Warning | 3 waters at 2.11–2.20 Å | Resolved (deleted, incl. active-site HOH843 with reason) |
| 1U19 | `incomplete_additives` | Warning | 2 free palmitate fragments missing O1/O2 | Resolved (deleted; palmitoyls built as Cys322/323 thioesters) |
| 6OIM | `chain_breaks` | Warning | Lys104 ⟶ Asp108 break (7.5 Å) | Resolved (residues 105–107 built) |

*(continued)*

| System | Finding code | Level | Reported condition | Outcome |
|---|---|---|---|---|
| 6OIM | `constant_volume_density_low` | Warning | Starting-box density 0.967 g $mL^{-1}$ | Resolved (job rewritten with explicit pressure) |

**Totals across the preparations:** 27 distinct findings (24 warnings, 3 informational, 0 errors): 17 warnings resolved, by the preparation itself or by a decision of the agent stated in its report (the 1M17 loop, the 4EIY membrane box, the deleted 4WB5 water, the rewritten 6OIM job, the deleted 1U19 palmitate fragment), 6 accepted with written justification, 1 resolved in the bonded model and accepted in the non-bonded control. Ten follow-up messages reached the preparation agents after their first prompt; three carried scientific decisions (1STP/4DFR/1A28 stoichiometry, the 1A6N His64 tautomer and the 6OIM $Mg^{2+}$ model). These and the decisions taken during analysis are listed below.

**Decisions taken by the scientist.** What the agents reported or asked, and what the scientist decided, during preparation and analysis.

| System | What the agent reported or asked | The scientist's decision |
|---|---|---|
| 1STP, 4DFR, 1A28 | 1STP holds one subunit of the streptavidin tetramer and no tool rebuilds the assembly; 4DFR and 1A28 have two copies in the asymmetric unit | 1STP excluded; chain A only for 4DFR and 1A28 (the agent rebuilt both jobs) |
| 1A6N | PROPKA set the distal His64 to HID; the agent noted that His64 is often modeled as HIE in myoglobin and asked | HIE (the agent recorded the manual override). Neutron diffraction places the His64 hydrogen on Nε2 in oxymyoglobin at pH 8.4[1] but on Nδ1 in carbonmonoxymyoglobin at pH 5.7[2] and in metmyoglobin at pH 6.2[3]; in deoxymyoglobin the distal water is hydrogen-bonded to Nε2[4], and X-ray data cannot assign the tautomer[1]. |
| 6OIM | With a brief that named only the biological components, the agent kept the default $Mg^{2+}$ model (Li–Merz 12-6), being unsure how the $r^{-4}$ term of the 12-6-4 model acts on GDP and the OpenFF-typed adduct, and offered 12-6-4 as an option | 12-6-4, the model that better describes $Mg^{2+}$ bound to phosphates and already used for PKA, with the explanation that the $r^{-4}$ term acts on every atom according to its element and hybridization (the agent rebuilt the job) |
| 1ZAA, 3HS4 | With the first bonded parameters (RESP charges at the HF/6-31G* level), the run check reported the carbonyl oxygen of Ile67 within 2.5 Å of Zn203 in 35% and 23% of the frames of two Zif268 replicas | The zinc sites parameterized again, with B3LYP/6-31G* RESP charges and a revised bonded model; with these the contact lasts 0.5 ns in one replica (Fig. 3g); 1IRO was prepared in the same way |
| 1ZAA, 1IRO | The agent kept the QM equilibrium angles, forecasting that S–Zn–S would open to about 125°, and stopped when the QM of 1IRO could run only on the CPU, which its instructions did not allow | Both accepted after review |
| 1U19 | The agent deleted a mis-modeled mannose on Asn15 and placed this judgment call at the top of its report | The agent's choices reviewed; the job submitted unchanged |

*(continued)*

| System | What the agent reported or asked | The scientist's decision |
|---|---|---|
| Protein–ligand complexes (the decision applied to every MD system) | Automatic equilibration detection discarded 100–145 ns in several replicas for drifts of 0.1–0.4 Å | Every quantity reported for the analyzed window and for the whole production (Supplementary Data 3) |
| `1M17` | The modeled C-terminal tail reaches its periodic image in two replicas; the analysis agent judged the box too small for the extended tail | Runs kept and analyzed on the kinase core, with the contact reported; not rerun |
| `1M17` | The agent gave the DFG-in thresholds from memory and asked to have them checked | Thresholds checked in the Kincore paper[5] before citing; the agent's assignment stood |
| `4EIY` | An analysis report gave ff19SB as the protein force field of the run | Corrected to ff14SB from the run's own record; later briefs asked the agents to read the force fields of each run from its entry |
| `1U19` | The direct Schiff-base–Glu113 salt bridge opens late in two replicas | A follow-up analysis of the solvent around the Schiff base (Supplementary Section 4) |
| Alanine dipeptide | With umbrella windows of 2 ns the estimate was still rising at the end of the windows | Windows run again for 5 ns |
| Alanine dipeptide | In the 5-ns windows each window starts with a transient that lowers the estimate by about 0.25 kcal mol$^{-1}$; the agent gave the result with and without it and left the choice to the scientist | The first 1.25 ns of each window excluded (1.61 ± 0.05 kcal mol$^{-1}$) |
| TYK2 | — | The blind test designed: receptor and posed ligands under anonymous names, a budget of about 1,000 GPU-hours, no lookup of the compounds |

**Supplementary Table 5 | Quantitative summary of the validation systems.** Summary of the properties in the analyzed windows: means ± s.d. across replicas, except per-frame s.d. for the single replica of `4EIY` and for the NH···S count of `1IRO`, and block or bootstrap errors for alanine dipeptide. The analysis agents placed each replica's window from the convergence analysis; at the scientist's request every value is also given over the whole production (Supplementary Data 3). Full per-replica values (both equilibrated window and whole production), thermodynamic checks, metal-site angles and lipid order parameters are provided in `Supplementary_Data_3_validation_results_per_replica.csv` and `Supplementary_Data_4_lipid_order_parameters.csv`.

| System (PDB, runs) | Key structural / dynamical observables (equilibrated window) | Reference / benchmark & notable replica events |
|---|---|---|
| **Soluble protein & protein–ligand complexes** | | |
| Ubiquitin (`1UBQ`, 5 × 1 µs) | Core backbone RMSD (1–70) 0.80 ± 0.03 Å; native contacts $Q$ 0.951 ± 0.007; core $R_g$ 11.15 ± 0.03 Å; Cα B-factor correlation $r = 0.755$, $\rho = 0.606$ | Crystal `1UBQ` core $R_g$ 11.06 Å; Daura cluster 1 = 91.8 ± 4.0%; β1–β2 loop visits third sub-state in 85% of r3 (4–18% in others) |

*(continued)*

| System (PDB, runs) | Key structural / dynamical observables (equilibrated window) | Reference / benchmark & notable replica events |
|---|---|---|
| Progesterone receptor (1A28, 3 × 300 ns) | Backbone RMSD 1.30 ± 0.21 Å; progesterone RMSD 0.55 ± 0.02 Å (100% < 2 Å); Arg766–O3 H-bond 30 ± 26% | Chain A only (scientist decision); Arg766–O3 breaks at ~95 ns in r2 while pose remains locked; water-mediated 3-keto contacts in r1 |
| DHFR–methotrexate (4DFR, 3 × 300 ns) | Backbone RMSD 1.31 ± 0.15 Å; pteridine–benzoyl core RMSD 0.78 ± 0.31 Å; Asp27 pair 100%, Ile5 O 99.4 ± 0.7% | Chain A only (scientist decision); Met20 loop stays occluded in r3 (92.1%), never closed (< 1% vs 1RX2[6]); r1 glutamate tail rearranges at 12–18 ns (whole-ligand RMSD 3.26 Å) |
| HIV-1 protease–indinavir (1HSG, 3 × 300 ns) | Backbone RMSD 1.11 ± 0.14 Å; indinavir RMSD 1.38 ± 0.87 Å; O2⟶Asp25′ OD2 97.7–98.5% (r1, r3); flap water bridge 94.0 ± 6.4% | Flaps stay closed (Ile50–Ile50′ Cα ≤ 9.96 Å); reversible pyridylmethyl excursion out of S3 in r2 (130–270 ns, returns) |
| Estrogen receptor α–4-OHT (3ERT, 3 × 300 ns) | Core backbone RMSD 1.86 ± 0.32 Å; 4-OHT RMSD 1.30 ± 0.18 Å (96–100% < 2 Å); Helix 12 RMSD 1.27–1.45 Å | Antagonist H12 conformation preserved; phenol–Glu353 direct (61.3 ± 32.9%) and water-bridged (57.5 ± 25.8%); Asp351 salt bridge intermittent (10–35%) |
| EGFR kinase–erlotinib (1M17, 3 × 300 ns) | Core backbone RMSD 1.68 ± 0.31 Å; erlotinib RMSD 1.47 ± 0.14 Å; hinge Met769 N⟶N1 79.7 ± 3.6%; DFG-in 100% | Kincore DFG-in criteria[5]; Lys721–Glu738 lost in r2 (5.2%); post-run check flagged the drift; the analysis agent measured C-terminal tail contacts with the periodic image in r1 (2.44 Å) and r3 (2.49 Å); runs kept and the contact reported by the scientist's decision |
| **Lipid bilayers & membrane receptors** | | |
| Pure POPC (3 × 1 μs, 303.15 K) | Area/lipid 59.76 ± 0.08 Å$^2$; $D_{PP}$ 40.75 ± 0.03 Å; $D_{HH}$ 39.97 ± 0.18 Å; $-S_{CD}$ plateaus 0.231 (*sn*-1) / 0.217 (*sn*-2); $D$ 1.66 ± 0.14 μm$^2$ s$^{-1}$ | Area ~7% below the experimental ~64 Å$^2$[27], flagged unprompted by the analysis agent; symmetric leaflets, zero flip-flop |
| POPC–cholesterol 29.7 mol% (3 × 1 μs) | Area/mol 46.44 ± 0.16 Å$^2$ (−22%); Voronoi POPC 50.38 ± 0.23, CHL1 37.13 ± 0.02 Å$^2$; $D_{PP}$ 45.13 ± 0.12 Å (+4.4 Å); $-S_{CD}$ +45% / +37%; CHL1 tilt 14.66 ± 0.17° | Reproduces cholesterol condensation, thickening, acyl ordering[8] and ~50% slower POPC diffusion (0.88 ± 0.15 μm$^2$ s$^{-1}$); r2 area relaxes until 265 ns |
| A2A receptor in POPC (4EIY, 1 × 500 ns) | TM backbone RMSD 1.47 ± 0.10 Å (98.8% helical); ZM241385 RMSD 1.55 ± 0.22 Å; Asn253 H-bonds 95.2%; Phe168 stacking 3.89 ± 0.26 Å | Allosteric $Na^+$ retained on Asp52 (6.00 ± 0.58 shell atoms)[9]; condensed local Voronoi APL 54.88 ± 0.63 Å$^2$, $D_{PP}$ 41.06 ± 0.36 Å |
| Rhodopsin asymmetric disc (1U19, 3 × 200 ns) | TM RMSD 0.73 ± 0.06 Å; ionic lock 4.65 ± 0.13 Å; retinal RMSD 0.56 ± 0.03 Å, C11=C12 −6.7 ± 0.9°; box area/mol 58.88 ± 0.27 Å$^2$; $D_{PP}$ 44.79 ± 0.14 Å | All 33 PS retained in cytoplasmic leaflet (0 flip-flop); glycans intradiscal, palmitoyls cytoplasmic; direct NZ–Glu113 salt bridge formed for ~100–140 ns, then intermittent and broken over the last ~30–50 ns of r1/r2, bridged by crystal waters (follow-up analysis requested by the scientist) |
| **Metalloproteins, heme, phosphorylated & covalent complexes** | | |

*(continued)*

| System (PDB, runs) | Key structural / dynamical observables (equilibrated window) | Reference / benchmark & notable replica events |
|---|---|---|
| Zif268–DNA (1ZAA, 2 × 3 × 200 ns) | **Bonded:** CN 4.00 at all 3 sites, 0 of 36 donor–replica pairs lost, Zn–N 1.98–2.03 Å, Zn–S 2.23–2.28 Å, backbone RMSD 1.56 ± 0.23 Å. **Non-bonded:** 9/18 His donor–replica pairs lost, Zn–S collapses to 1.84–1.87 Å, backbone RMSD 2.46 ± 0.83 Å | Crystal Zn–S 2.15–2.44 Å[10]; all 10 DNA base pairs intact (≥ 98%) in both; with the first bonded parameters Ile67 O came within 2.5 Å of Zn203 in 35% and 23% of the frames of two replicas; after the scientist had the sites parameterized again, with B3LYP/6-31G* RESP charges and a revised bonded model, 0.5 ns in one replica |
| Carbonic anhydrase II (3HS4, 2 × 3 × 200 ns) | **Bonded:** 0 donors lost, Zn–N 1.93–2.05 Å, AZM RMSD 0.93 ± 0.02 Å, Thr199 H-bonds 91.9% / 70.5%. **Non-bonded:** His94 lost before production, Glu106 & AZM O2 bind at 1.69–1.73 Å, AZM RMSD 3.14 ± 0.80 Å | Bonded site preserves tetrahedral geometry (99–116°) and Thr199 network; non-bonded control destroys active-site geometry and collapses Thr199 contacts (< 5%) |
| Oxidized rubredoxin (1IRO, 2 × 3 × 200 ns) | **Bonded:** CN 4.00, Fe–S 2.27–2.31 Å (within 0.06 Å of crystal), NH···S H-bonds 4.21 ± 0.97/frame, backbone RMSD 0.88 ± 0.03 Å. **Non-bonded:** CN 6.00 (Val8 O & Leu41 O bind at 2.07–2.08 Å), Fe–S 2.13–2.17 Å, S9–Fe–S42 163.1° | Non-bonded Li–Merz $Fe^{3+}$ collapses tetrahedral $Fe(Cys)_4$ into octahedral $Fe(S)_4(O)_2$ and loses Cys9⟶S6 & Cys42⟶S39 NH···S bonds |
| Deoxymyoglobin (1A6N, 3 × 200 ns) | Backbone RMSD 1.01 ± 0.02 Å; porphyrin core 0.58 ± 0.04 Å; Fe–His93 2.044 ± 0.002 Å; Fe out-of-plane +0.069 ± 0.002 Å; His64 H-bond to distal HOH1001 74.9 ± 4.0% | Crystal Fe–His93 2.14 Å, doming +0.364 Å (Amber heme model flattens Fe in-plane during prep); His64 set to HIE by scientist |
| PKA–PKI–ATP–2 $Mg^{2+}$ (4WB5, 3 × 200 ns) | Kinase RMSD 1.23 ± 0.11 Å; ATP RMSD 0.78 ± 0.09 Å; Mg402/Mg403 CN 5.99 ± 0.02 / 6.03 ± 0.03; Lys72–Glu91 & PKI Arg salt bridges 100%; Lys189–pThr197 89.9 ± 11.7% | ATP bridging oxygen O3B leaves Mg402 for good in all 3 replicas (24–59 ns by the analysis's persistence rule) and a bulk water takes its place; Mg coordination was on the preparation agent's watch-list |
| KRAS G12C–sotorasib (6OIM, 3 × 200 ns) | Backbone RMSD 0.96 ± 0.16 Å; Cys12 Sγ–C25 1.869 ± 0.001 Å (max 2.02 Å); sotorasib RMSD 0.86 ± 0.03 Å; GDP RMSD 0.41 ± 0.01 Å; $Mg^{2+}$ CN 6.00 (no exchange) | Scientist chose the 12-6-4 $Mg^{2+}$ model (the agent had kept the default and offered 12-6-4); Switch II shifts ~2 Å in r2 (62–66 ns) without perturbing sotorasib or its His95/Tyr96/Gln99 contacts |
| **Enhanced sampling** | | |
| Alanine dipeptide (300 K, 4 protocols) | $\Delta G(\phi^+ - \phi^-)$: plain MD 1.54 ± 0.24 (500 ns), WTMetaD 1.70 ± 0.03 (100 ns), umbrella WHAM 1.61 ± 0.05 kcal $mol^{-1}$ (36 × 5 ns, first 1.25 ns discarded); barrier near $\phi = 0°$: 5.47–5.73 kcal $mol^{-1}$ across WTMetaD, WHAM and SMD | SMD profile matches WHAM within 0.09 kcal $mol^{-1}$ over −63° to +38°; umbrella window overlap 0.27–0.52; transient in the first 1.25 ns of each window flagged by the agent and excluded by the scientist |

**Supplementary Table 6 | Quantum-chemical models of the bonded metal sites.** The bonded metal sites of the validation, as Molpylot recorded them when it parameterized each site: the donors, the small model from whose Hessian the Seminario force constants were derived, and the capped large model to whose electrostatic potential the RESP charges were fitted. Both models of a site carry the same net charge, the metal's oxidation state plus the force-field charge of each coordinating residue (thiolate cysteines −1, histidines 0, acetazolamide as the sulfonamide anion −1), and the same spin multiplicity. All calculations used B3LYP/6-31G* (Cartesian d functions) in PySCF: a constrained optimization of the small model, with the link hydrogens and the atoms carrying them held in place; its analytical Hessian, with the bond lengths and angles of the optimized model as equilibrium values; and a two-stage RESP fit on four shells at 1.4–2.0 times the van der Waals radii, with the ACE/NME caps and the backbone N, H, C and O held at the force field's values and the metal restrained toward its formal charge. In the small model the side chains are cut between Cα and Cβ, the Cα replaced by a link hydrogen; in the large model the coordinating residues are whole and their neighbors become ACE/NME caps. Molpylot takes as donors the atoms bonded to the metal in the structure or within element-pair cutoffs (Fe–S 2.65 Å, Zn–N 2.45 Å, Zn–S 2.75 Å); here they are exactly the metal–donor bonds recorded in the deposited entries. Distances (Å) are those of the prepared structures, which keep the crystal coordination. Residue numbers are those of the deposited entries. The first parameterization of the zinc sites, before the scientist had them parameterized again (Supplementary Table 4), used the same donors and the same models, with RESP charges fitted at the HF/6-31G* level and an earlier form of the bonded terms.

| Site | Metal state | Donors (distance, Å) | Small model (Seminario Hessian) | Large model (RESP) | Net charge, multiplicity |
|---|---|---|---|---|---|
| Zif268–DNA (1ZAA) Zn201 | Zn(II), closed shell | Cys7 SG 2.23, Cys12 SG 2.44, His25 NE2 2.14, His29 NE2 1.88 | Zn, Cys and His side chains from Cβ; 35 atoms (4 link H) | Zn, Cys7, Cys12, His25, His29 whole; caps from residues 6, 8, 11, 13, 24, 26, 28, 30; 103 atoms | 0, singlet |
| Zif268–DNA (1ZAA) Zn202 | Zn(II), closed shell | Cys37 SG 2.34, Cys40 SG 2.21, His53 NE2 2.00, His57 NE2 2.01 | Zn, Cys and His side chains from Cβ; 35 atoms (4 link H) | Zn, Cys37, Cys40, His53, His57 whole; caps from residues 36, 38, 39, 41, 52, 54, 56, 58; 103 atoms | 0, singlet |
| Zif268–DNA (1ZAA) Zn203 | Zn(II), closed shell | Cys65 SG 2.23, Cys68 SG 2.15, His81 NE2 2.12, His85 NE2 1.98 | Zn, Cys and His side chains from Cβ; 35 atoms (4 link H) | Zn, Cys65, Cys68, His81, His85 whole; caps from residues 64, 66, 67, 69, 80, 82, 84, 86; 103 atoms | 0, singlet |
| Carbonic anhydrase II–acetazolamide (3HS4) Zn301 | Zn(II), closed shell | His94 NE2 1.99, His96 NE2 1.99, His119 ND1 2.01, acetazolamide N1 1.94 (a) | Zn, His side chains from Cβ, whole acetazolamide; 55 atoms (3 link H) | Zn, His94, His96, His119, acetazolamide whole; caps from residues 93, 95, 97, 118, 120; 101 atoms | +1, singlet |
| Oxidized rubredoxin (1IRO) Fe55 (b) | Fe(III), high spin, 2S = 5 (c) | Cys6 SG 2.29, Cys9 SG 2.25, Cys39 SG 2.28, Cys42 SG 2.23 | Fe, Cys side chains from Cβ; 21 atoms (4 link H) | Fe, Cys6, Cys9, Cys39, Cys42 whole; caps from residues 5, 7, 8, 10, 38, 40, 41, 43; 89 atoms | −1, sextet |

(a) The sulfonyl S1 of acetazolamide, 3.05 Å from the zinc, was reported as a candidate beyond the Zn–S cutoff and not bound.

(b) The Hessian and the electrostatic potential of this site were computed on the CPU (Supplementary Table 4); those of the zinc sites on the GPU.

(c) Given in the parameterization call; unrestricted calculations, with $\langle S^2 \rangle = 8.76$ (8.75 for S = 5/2). The states of the zinc ions were assigned from the residue name.

**Supplementary Table 7 | TYK2 relative binding free-energy campaign statistics.** Comparison of the published 22-edge reference network and the blind 24-edge agent-planned network (16 ligands[11]; OpenFE 1.13 hybrid-topology replica exchange, 11 $\lambda$ windows × 5 ns × 3 repeats/leg, OpenFF Sage 2.3.0/NAGL) against published OpenFE benchmark results recomputed from the deposited per-ligand values of Baumann et al.[12]. Values in kcal $mol^{-1}$ with 95% bootstrap confidence intervals. Complete per-ligand $\Delta G$, per-edge $\Delta\Delta G$ and cycle-closure tables are provided in `Supplementary_Data_5_tyk2_fep_ligands.csv`, `Supplementary_Data_6_tyk2_fep_edges.csv` and `Supplementary_Data_7_tyk2_fep_cycles.csv`.

| Metric | Reference network (22 edges) | Agent-planned network (24 edges) | Published OpenFE[12] |
|---|---|---|---|
| Ligands / edges / cycles | 16 / 22 / 7 | 16 / 24 / 9 | 16 / — / — |
| Completed edges | 22 / 22 | 24 / 24 | — |
| $\Delta G$ RMSE | 0.60 [0.43, 0.76] | 0.59 [0.43, 0.73] | 0.59 |
| $\Delta G$ MUE | 0.51 [0.36, 0.67] | 0.52 [0.39, 0.65] | — |
| $\Delta G$ $R^2$ | 0.79 [0.61, 0.92] | 0.78 [0.60, 0.91] | 0.80 |
| $\Delta G$ Kendall $\tau$ | 0.65 [0.39, 0.86] | 0.70 [0.46, 0.89] | 0.68 |
| $\Delta\Delta G$ RMSE / MUE | 0.81 / 0.72 | 0.94 / 0.79 | — |
| $\Delta\Delta G$ $R^2$ / $\tau$ | 0.50 / 0.40 | 0.32 / 0.32 | — |
| Largest cycle closure | +0.97 ± 0.48 (`ejm_42→50→49→31`) | +0.61 ± 0.52 (`lig07→02→14→16`) | — |
| Lowest neighboring-window overlap | 0.052 (`ejm_31~ejm_44`) | 0.071 (`lig02~lig07`) | — |
| Worst repeat spread vs per-repeat MBAR error | `ejm_55~ejm_54`: 0.29 vs 0.23 | `lig08~lig11`: 0.60 vs 0.31 | — |

### Supplementary Section 1 | Soluble protein and protein–ligand complexes

**Common protocol.** Ubiquitin (`1UBQ`) and the five protein–ligand complexes (`1A28`, `4DFR`, `1HSG`, `3ERT`, `1M17`) were prepared by tool-confined agents with Molpylot's soluble defaults: Amber ff19SB[13], OPC water[14], rhombic dodecahedron with 10 Å padding, 0.15 M NaCl, PROPKA protonation at pH 7, crystallographic waters retained within 5 Å of heteroatoms, OpenFF Sage 2.3.0 with NAGL charges for ligands[15], and the 10-stage Roe–Brooks equilibration protocol[16]. Production lengths were 5 × 1 µs (`1UBQ`) and 3 × 300 ns (each complex). The analysis agents placed each replica's analyzed window from the convergence analysis of the backbone and ligand RMSD, departing from the detector where they judged it necessary; at the scientist's request every quantity is also reported over the whole production (Supplementary Data 3).

**Ubiquitin (`1UBQ`).** No pre-execution warnings arose; before job submission, the agent caught that the default SLURM header pointed to a debug partition unable to host 1 µs runs and requested the production header. Across 100–1000 ns (25,837 atoms), the core fold (residues 1–70) remains stable (RMSD 0.80 ± 0.03 Å, $Q = 0.951 \pm 0.007$), and per-residue fluctuations correlate with crystallographic Cα B factors ($r = 0.755$, $\rho = 0.606$). Joint PCA on residues

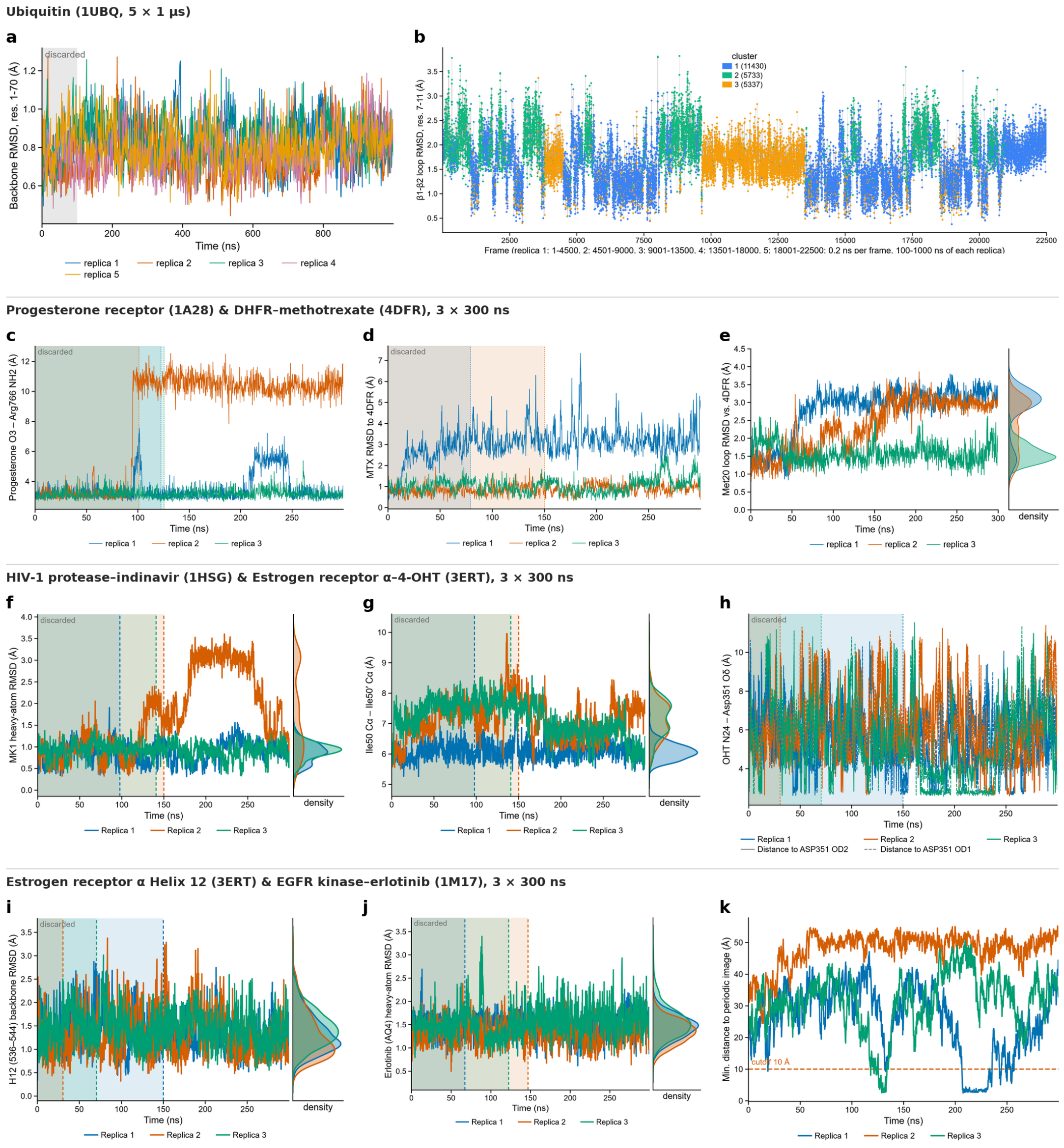


**Supplementary Fig. 1 | Soluble protein and protein–ligand complex validation (1UBQ, 1A28, 4DFR, 1HSG, 3ERT and 1M17).** **a**,**b**, Ubiquitin (`1UBQ`, 5 × 1 µs): core backbone RMSD (residues 1–70; 0–100 ns discard shaded) (**a**) and β1–β2 loop (residues 7–11) RMSD across all five concatenated replicas colored by *k*-means sub-state, highlighting replica 3's occupancy of state 3 (**b**). **c**, Progesterone receptor (`1A28`, 3 × 300 ns): progesterone O3–Arg766 Nη2 distance showing loss at ~95 ns in replica 2 and transient excursion in replica 1. **d**,**e**, DHFR–methotrexate (`4DFR`, 3 × 300 ns): whole-ligand heavy-atom RMSD showing the replica 1 glutamate-tail rearrangement (**d**) and Met20 loop backbone RMSD relative to occluded `4DFR` (**e**). **f**,**g**, HIV-1 protease–indinavir (`1HSG`, 3 × 300 ns): indinavir heavy-atom RMSD showing the reversible replica 2 pyridylmethyl excursion (130–270 ns) (**f**) and Ile50–Ile50′ Cα flap distance confirming closed flaps (**g**). **h**,**i**, Estrogen receptor α–4-hydroxytamoxifen (`3ERT`, 3 × 300 ns): OHT N24–Asp351 Oδ1/Oδ2 salt-bridge distances (**h**) and Helix 12 (residues 536–544) backbone RMSD confirming retention of the antagonist conformation (**i**). **j**,**k**, EGFR kinase–erlotinib (`1M17`, 3 × 300 ns): erlotinib heavy-atom RMSD (**j**) and minimum protein-to-periodic-image distance with the 10 Å threshold dashed, flagging the C-terminal tail image contacts in replicas 1 and 3 (**k**). Shaded regions denote per-replica discarded equilibration windows.

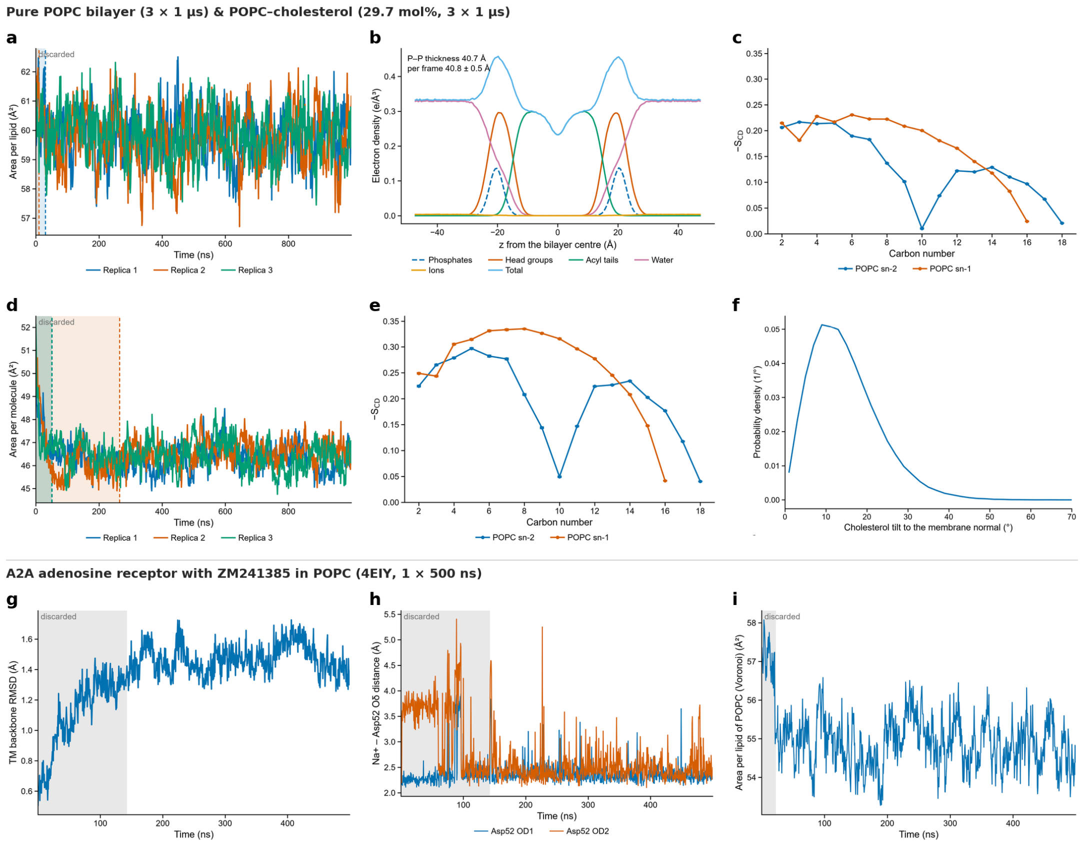


**Supplementary Fig. 2 | Lipid bilayers (POPC, POPC–cholesterol) and A2A adenosine receptor (4EIY). a–c**, Pure POPC bilayer (3 × 1 µs at 303.15 K): area per lipid time series (**a**), pooled electron-density profile across the bilayer normal (**b**), and deuterium order parameters ($-S_{CD}$) of *sn*-1 and *sn*-2 acyl chains (**c**). **d–f**, POPC–cholesterol bilayer (29.7 mol% cholesterol, 3 × 1 µs at 303.15 K): box area per molecule showing slower relaxation of replica 2 until 265 ns (**d**), elevated POPC $-S_{CD}$ order parameters on the same vertical scale as **c** (**e**), and cholesterol C3⟶C17 tilt distribution relative to the membrane normal (**f**). **g–i**, A2A adenosine receptor–ZM241385 embedded in POPC (4EIY, 1 × 500 ns): transmembrane-bundle backbone RMSD (**g**), allosteric $Na^+$–Asp52 Oδ1/Oδ2 coordination distances showing transition to bidentate binding near 100 ns (**h**), and Voronoi area per POPC molecule around the receptor (**i**).

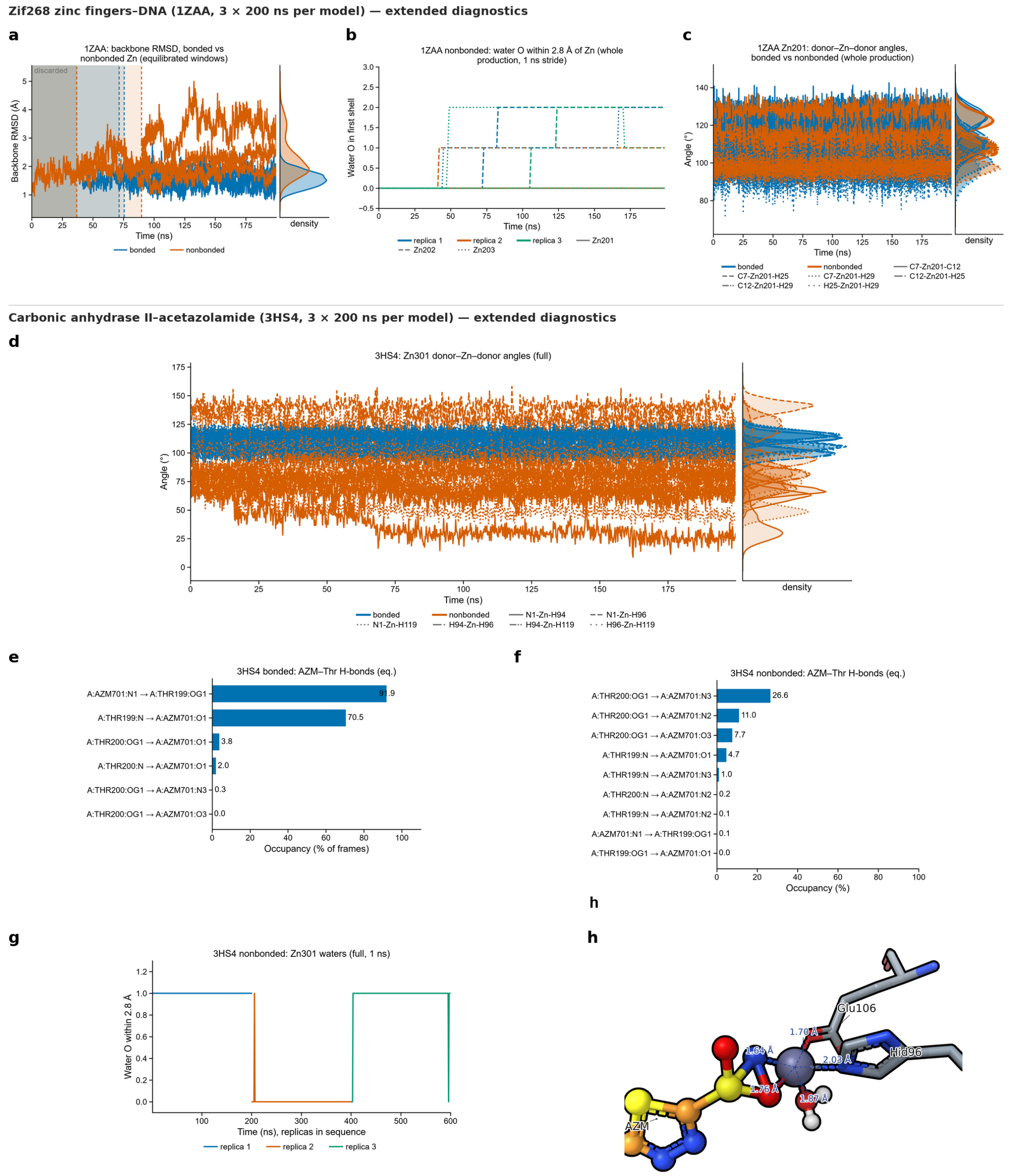


**Supplementary Fig. 3 | Extended zinc-site diagnostics for Zif268 (1ZAA) and carbonic anhydrase II (3HS4).** Complementing Fig. 3d–g; bonded B3LYP/6-31G* model (blue) versus non-bonded Li–Merz $Zn^{2+}$ control (orange) over 3 × 200 ns per model. **a–c**, Zif268–DNA (1ZAA): protein backbone RMSD in the post-equilibration windows (**a**), number of water oxygens entering within 2.8 Å of each zinc ion in the non-bonded control (**b**), and donor–Zn–donor angles at the intact Zn201 site (**c**). **d–h**, Carbonic anhydrase II–acetazolamide (3HS4): donor–Zn–donor angles across full production showing distortion of the non-bonded tetrahedron (**d**), acetazolamide–Thr199/Thr200 hydrogen-bond occupancies in the bonded (**e**) and non-bonded (**f**) models, first-shell water intrusion in the non-bonded control across concatenated replicas (**g**), and 3D snapshot of the collapsed non-bonded zinc site (replica 1, 149.1 ns) showing Glu106 Oε1, acetazolamide O2 and water coordination (**h**).

1–70 resolves three β1–β2 loop sub-states (Leu8–Lys11); replica 3 spends 85.0% of its frames in state 3 (versus 4.4–18.2% in other replicas), which caused automated convergence detection to flag 439 ns for replica 3, an effect the analysis agent correctly diagnosed as a loop switch rather than fold relaxation, retaining the common 100 ns discard (Supplementary Fig. 1a,b).

**Scientist decision: stoichiometry (`1STP`, `4DFR`, `1A28`).** While preparing seven candidate complexes, the agent halted on monomeric streptavidin (`1STP`, whose biological tetramer no tool could assemble) and flagged the two crystal copies in `4DFR` and `1A28`. The scientist excluded `1STP` and selected chain A for `4DFR` and `1A28`, after which the agent rebuilt both jobs.

**Complex-specific findings.**

- `1A28` **(progesterone receptor).** Five incomplete side chains were rebuilt automatically. Progesterone remains locked (0.55 ± 0.02 Å, 100% < 2 Å) even when the direct Arg766–O3 hydrogen bond breaks near 95 ns in replica 2 (Supplementary Fig. 1c); in replica 1 the 3-keto group is also held by exchanging water bridges to Arg766, Met759 and Gln725.
- `4DFR` **(DHFR–methotrexate).** Pre-checks flagged three strained C–O bonds (relaxed by minimization), nine incomplete residues (rebuilt) and the native *cis* Gly95–Gly96 peptide (kept). The agent kept methotrexate at −1 (N1 protonated) to match the Asp27 salt bridge and kept one chain-B crystal water within 5 Å of chain A's ligand. While the pteridine–benzoyl core stays bound (0.78 ± 0.31 Å; Asp27 100%), replica 1's glutamate tail rearranges at 12–18 ns (breaking Arg57 and forming a Lys32 contact). To classify the Met20 loop, the analysis agent found that hydrogen-bond markers flickered across replicas, fetched the closed reference `1RX2`[6], and applied a 2.0 Å backbone RMSD criterion: no replica adopted the closed state (< 1% of frames), consistent with the absence of NADPH (Supplementary Fig. 1d,e).
- `1HSG` **(HIV-1 protease–indinavir).** PROPKA and H-bond optimization assigned the canonical monoprotonated catalytic dyad (Asp25 protonated, Asp25′ deprotonated). Flaps remain closed (Ile50–Ile50′ Cα ≤ 9.96 Å) with the flap-water site continuously occupied (94.0 ± 6.4%, exchanging among crystal and bulk waters across replicas). In replica 2, the pyridylmethyl group reversibly leaves the S3 subsite between 130 and 270 ns and returns (Supplementary Fig. 1f,g).
- `3ERT` **(estrogen receptor α–4-OHT).** Twelve incomplete residues were rebuilt and backbone-less Pro552 removed. 4-OHT (protonated amine) and the antagonist Helix 12 conformation remain stable (1.30 ± 0.18 Å and 1.27–1.45 Å). Inspecting the preparation stage structures, the analysis agent established that the borderline Asp351–N24 salt bridge (3.90 Å in crystal) opens during equilibration in replicas 1–2 and reforms intermittently across replicas (10–35%; Supplementary Fig. 1h,i).
- `1M17` **(EGFR kinase–erlotinib).** The agent closed the His964⟶Leu977 chain break by modeling residues 965–976 (`LPSPTDSNFYRA`). Erlotinib (1.47 ± 0.14 Å), its Met769 hinge H-bond (79.7 ± 3.6%) and the DFG-in state (100% by the Kincore criteria[5], which the agent recalled from memory and the scientist verified in the paper) are maintained across replicas, while Lys721–Glu738 breaks in replica 2 (5.2%). Post-run checks flagged large structural drift, and the analysis agent measured periodic-image contacts of the extended C-terminal tail in replicas 1 (2.44 Å) and 3 (2.49 Å; Pro995 O contacting Ser728 Oγ of the image), whereas replica 2 stayed ≥ 24.4 Å away (Supplementary Fig. 1j,k); the analysis

agent concluded that the box is too small for the extended tail. **Scientist decision:** keep the runs, analyze the kinase core and report the contact, rather than rerun the system.

#### Supplementary Section 2 | Lipid bilayers and A2A adenosine receptor

**Pure POPC and POPC–cholesterol bilayers (3 × 1 µs each, 303.15 K).** Built with Molpylot's membrane builder using Lipid21[17], TIP3P and Joung–Cheatham KCl (98 POPC/leaflet; or 78 POPC + 33 CHL1/leaflet, 29.7 mol%) with semi-isotropic Monte Carlo barostatting. Pure POPC equilibrated within 0–30 ns to an area per lipid of 59.76 ± 0.08 Å$^2$ (~7% below the experimental ~64 Å$^2$ [7], an offset flagged unprompted by the analysis agent), $D_{PP} = 40.75 \pm 0.03$ Å and lateral diffusion 1.66 ± 0.14 µm$^2$ s$^{-1}$. Adding 29.7 mol% cholesterol slowed area relaxation (requiring up to 265 ns in replica 2, detected automatically by the membrane-area check) and reproduced the condensing effect: area per molecule fell 22% to 46.44 ± 0.16 Å$^2$ (Voronoi POPC 50.38 ± 0.23, CHL1 37.13 ± 0.02 Å$^2$), $D_{PP}$ thickened by 4.4 Å to 45.13 ± 0.12 Å, *sn*-1/*sn*-2 $-S_{CD}$ plateaus rose by 45% and 37%[8], POPC diffusion nearly halved (0.88 ± 0.15 µm$^2$ s$^{-1}$), and cholesterol tilted at 14.66 ± 0.17° (Supplementary Fig. 2a–f; per-carbon $-S_{CD}$ in `Supplementary_Data_4_lipid_order_parameters.csv`). No flip-flop or core water penetration occurred in either system.

**A2A adenosine receptor with ZM241385 in POPC (`4EIY`, 1 × 500 ns).** Pre-checks raised four warnings, all resolved autonomously (BRIL loop 1043–1057 built, 7 incomplete residues rebuilt, ZM241385 parameterized, crystal cell replaced by membrane box). The agent selected the membrane preset (ff14SB, TIP3P, Lipid21), retained crystal waters (`waters="keep"`) and removed three crystal cholesterols lacking templates, documenting both departures. An analysis report that gave ff19SB as the protein force field of the run was corrected by the scientist from the run's own record. Over 500 ns (equilibration cutoffs 142.5 ns for protein/ligand, 22.6 ns for bilayer), the transmembrane bundle held its fold (RMSD 1.47 ± 0.10 Å, 98.8% helical), ZM241385 maintained its Asn253 H-bonds (95.2%) and Phe168 π-stacking (3.89 ± 0.26 Å), and the allosteric $Na^+$ ion[9] remained bound to Asp52 (switching from monodentate to bidentate near 100 ns; 6.00 ± 0.58 shell atoms; Supplementary Fig. 2g–i). Surrounding POPC condensed to 54.88 ± 0.63 Å$^2$ (upper 56.7, lower 53.1 Å$^2$) with $D_{PP} = 41.06 \pm 0.36$ Å.

#### Supplementary Section 3 | Metalloproteins, heme, phosphorylated and covalent complexes

**Bonded versus non-bonded metal sites (`1ZAA`, `3HS4`, `1IR0`; 2 × 3 × 200 ns each).** Prepared by one agent, comparing the quantum-derived bonded MCPB model[18,19] (B3LYP/6-31G* Hessian and RESP charges in PySCF; Supplementary Table 6) against Li–Merz non-bonded ion controls in ff19SB/OPC. The scientist designed the comparison and, after the run check on the first bonded model of Zif268 (RESP charges at the HF/6-31G* level) reported the carbonyl oxygen of Ile67 within 2.5 Å of Zn203, had the zinc sites parameterized again, with B3LYP/6-31G* RESP charges and a revised bonded model (Fig. 3g); the runs analyzed here use these parameters, and `1IR0` was prepared in the same way. In the trajectories, donors were defined within 2.8 Å (lost if absent in > 10% of frames).

- `1ZAA` **(Zif268–DNA, three $Zn(Cys)_2(His)_2$ sites)** and `3HS4` **(carbonic anhydrase II–acetazolamide).** Core results are presented in Fig. 3d–g and the main text. Complementary diagnostics (Supplementary Fig. 3) confirm that the bonded model preserves backbone stability, tetrahedral donor angles (93–125° in `1ZAA`, 99–116° in `3HS4`) and the acetazolamide–Thr199 hydrogen-bond network (91.9% and 70.5%), whereas non-bonded controls suffer heavy water intrusion, severe angular distortion (down to 55.5° in `3HS4`) and Thr199 H-bond collapse (< 5%). In `1ZAA`, the preparation agent warned in advance that using QM equilibrium angles would widen S–Zn–S from 111–116° (crystal) to ~125°, which MD confirmed (119.7–124.1°). In `3HS4`, the agent retained the sulfonamide anion (favored by 8.5 kcal $mol^{-1}$ in the binding site) and verified that sulfonamide O2 remained at 3.06 Å after minimization before exporting jobs (in MD, O2 flickers within 2.8 Å in 44–46% of frames, mean 2.69 Å).
- `1IRO` **(oxidized rubredoxin, high-spin Fe(III) $Fe(Cys)_4$, 2S = 5).** The agent verified the sextet spin state ($\langle S^2 \rangle = 8.76$, fitted Fe charge $+0.97\,e$, force constants 84–89 kcal $mol^{-1}$ $Å^{-2}$). Over 3 × 200 ns, the bonded site retains all four Fe–S bonds within 0.06 Å of the 1.1 Å crystal structure (2.27–2.31 Å, CN 4.00) and preserves the protective NH···Sγ H-bond network (4.21 ± 0.97/frame). By contrast, the non-bonded Li–Merz 12-6 $Fe^{3+}$ ion recruits the backbone carbonyl oxygens of Val8 and Leu41 (2.07–2.08 Å), collapsing into an octahedral $Fe(S)_4(O)_2$ complex (*trans* O–Fe–S 173–174°, S9–Fe–S42 163.1°) and destroying the Cys9⟶S6 and Cys42⟶S39 hydrogen bonds (Extended Data Fig. 4).

**Deoxymyoglobin (`1A6N`, 3 × 200 ns).** Prepared with the Amber high-spin Fe(II) heme model bonded to proximal His93. PROPKA assigned the distal His64 as HID; the agent noted that His64 is often modeled as HIE in myoglobin and asked. **Scientist decision:** HIE. Neutron diffraction places the His64 hydrogen on Nε2 in oxymyoglobin at pH 8.4[1] but on Nδ1 in carbonmonoxymyoglobin at pH 5.7[2] and in metmyoglobin at pH 6.2[3]; in deoxymyoglobin the distal water is hydrogen-bonded to Nε2[4], and X-ray data cannot assign the tautomer[1]. The agent rebuilt the system, recorded the manual override and corrected its own initial overstatement of a crystal H-bond in the HID state. The fold (1.01 ± 0.02 Å) and porphyrin core (0.58 ± 0.04 Å) remain stable, and distal crystal water HOH1001 stays in the pocket throughout all replicas (74.9 ± 4.0% H-bonded to His64 Nε2–H). As traced by the analysis agent, the standard Amber heme force field flattens Fe into the porphyrin plane already during minimization (+0.069 ± 0.002 Å vs +0.364 Å crystallographic doming; Extended Data Fig. 2a–c).

**PKA with PKI(5–24), ATP and two $Mg^{2+}$ ions (`4WB5`, 3 × 200 ns).** Three phosphorylated residues (pSer139, pThr197, pSer338) were kept as dianions with OpenFF Sage 2.3.0/NAGL side chains, $ATP^{4-}$ was drawn from the cofactor library[20], and the agent, whose brief asked it to choose the $Mg^{2+}$ model, queried `search_docs` and selected the Li–Merz 12-6-4 model for phosphate coordination over the default 12-6 compromise set. It also deleted a clashing active-site crystal water (HOH843, 2.20 Å from Asp166 Oδ2) with a geometric rationale and added Mg402–O3B coordination to its pre-run watch-list. In production (Extended Data Fig. 2d–f), the kinase (1.23 ± 0.11 Å), ATP (0.78 ± 0.09 Å), Lys72–Glu91 and PKI arginine salt bridges remain intact, and both $Mg^{2+}$ ions stay six-coordinate. In all three replicas the ATP bridging oxygen O3B leaves Mg402 for good and a bulk water takes its place until 200 ns (24.3–58.5 ns by the analysis's persistence rule; in replica 1 the ribose O3′ holds the site from 9.0 to 30.3 ns).

**KRAS G12C covalently bound to sotorasib (6OIM, 3 × 200 ns).** Prepared from an open brief naming only the biological components. The agent deleted expression residue Gly0, capped Lys169 with NME, built missing residues 105–107, selected neutral sotorasib (79% at pH 7.4), parameterized the Cys12–sotorasib adduct with OpenFF Sage 2.3.0/NAGL, and pulled $GDP^{3-}$ from the cofactor library. **Scientist decision.** The agent kept the default $Mg^{2+}$ model (Li–Merz 12-6, `cm`): it could not confirm how the $r^{-4}$ term of the 12-6-4 model acts on GDP and the OpenFF-typed adduct atoms, and offered 12-6-4 as an option. The scientist chose 12-6-4, the model that better describes $Mg^{2+}$ bound to phosphates and already used for PKA, and explained that the $r^{-4}$ term acts on every atom according to its element and hybridization; the agent rebuilt the job and rewrote it cleanly after a `constant_volume_density_low` check. Across 3 × 200 ns (Extended Data Fig. 2g–i), the Cys12 Sγ–C25 bond holds at 1.869 ± 0.001 Å (max 2.02 Å), sotorasib (0.86 ± 0.03 Å) and GDP (0.41 ± 0.01 Å) keep their poses and contacts, and $Mg^{2+}$ retains all six crystal ligands without exchange. Switch II shifts ~2 Å in replica 2 (62–66 ns) without perturbing the inhibitor.

#### Supplementary Section 4 | Asymmetric rhodopsin disc membrane (1U19)

Complementing Fig. 3a–c and the main text, this section records the technical choices made autonomously by the preparation agent from a biological brief alone (brief in Supplementary Note 2) and the extended water-network analysis of the Schiff base (Extended Data Fig. 3a–d).

**Autonomous preparation choices.** The agent paired ff14SB/TIP3P with Lipid21 and GLYCAM-06j[21] (citing TIP3P parameterization consistency), parameterized retinal–Lys296 (explicitly specifying the 11-*cis*, 15-*anti* protonated Schiff base SMILES so the pH-7 default would not choose the neutral imine) and palmitoyl-Cys322/323 with Sage 2.3.0/NAGL, protonated Asp83 and Glu122 from dark-state FTIR data[22] (matching PROPKA $pK_a \approx 9.2$), followed PROPKA on Glu181 ($pK_a = 7.7$, explicitly flagging its debated protonation state), deleted a mis-modeled α-mannose in an inverted chair on Asn15 (placing this judgment call at the top of its report for the scientist to approve; the scientist reviewed the agent's choices and submitted the job unchanged), and expanded the membrane padding to 20 Å (111,829 atoms, 132 lipids/leaflet across POPC, SDPC, POPE, SDPE, POPS, SDPS and cholesterol) with a 5 ns final equilibration stage. It verified leaflet orientation directly from built coordinates and noted that replicas share initial lipid placements because the builder uses a fixed random seed.

**Schiff-base water network and area convergence.** While the receptor remains inactive (TM RMSD 0.73 ± 0.06 Å, ionic lock 4.65 ± 0.13 Å, retinal 11-*cis* within −31° to +18°), the direct Schiff-base Nζ–Glu113 salt bridge (≤ 4 Å), formed for about 100–140 ns, turns intermittent and is broken over the last ~30–50 ns of replicas 1 and 2 (broken in 21.2 ± 3.6% of all frames and 58.8 ± 25.3% of the last 50 ns; Extended Data Fig. 3a). A follow-up solvent analysis, requested by the scientist, showed that Nζ–H retains a hydrogen-bond partner in 98.5–100% of broken frames: almost exclusively one of three crystallographic waters, forming a one-water bridge to Glu113 (33.9 ± 13.1% of the last 50 ns) or a two-water chain, with zero bulk-water or ion entry; opening coincides in time with Ser186 donating a hydrogen bond to Glu113. Post-run membrane-area checks flagged slow ±1% area fluctuations ($\tau \approx 10$ ns) and incomplete formal area convergence in two replicas after the initial 6–7% contraction (Extended Data Fig. 3b).

#### Supplementary Section 5 | Alanine dipeptide enhanced sampling and TYK2 RBFE diagnostics

**Alanine dipeptide (300 K; complementing Fig. 4 and Extended Data Fig. 3e–h).** Ace-Ala-NMe (3,170 atoms, ff19SB, OPC, 0.15 M NaCl) was run across four protocols: 500 ns plain MD (200 + 300 ns continuous extension, junction heavy-atom RMSD 0.41 Å), 100 ns well-tempered metadynamics on $(\phi, \psi)$ (bias factor 10, 4 fs with HMR), 36 umbrella windows × 5 ns along $\phi$ ($k = 50$, 100 ps pre-equilibration each) and 20 steered-MD pulls ($\phi = -110° \rightarrow +60°$, $30°\ \mathrm{ns}^{-1}$, $k = 200\ \mathrm{kcal\ mol^{-1}\ rad^{-2}}$). The analysis agent fixed basin boundaries once from the metadynamics saddles ($\phi^{+}$: $0° \leq \phi < 120°$, excluding the $\beta$ tail at 120–180°; C7ax lies on a saddle in OPC water and is merged into $\alpha_{\mathrm{L}}$). A TICA Markov model on plain MD (lag 0.5 ns) yielded mean first-passage times of 28.7 ns ($\phi^{-} \rightarrow \phi^{+}$) and 2.16 ns ($\phi^{+} \rightarrow \phi^{-}$), matching direct counting (24.5 and 1.8 ns over 38 transitions). For umbrella sampling, consecutive 1.25 ns blocks per window gave $\Delta G = 1.355$, 1.632, 1.651 and 1.544 $\mathrm{kcal\ mol^{-1}}$, revealing a systematic $-0.25\ \mathrm{kcal\ mol^{-1}}$ initial transient. The windows had been run for 5 ns at the scientist's request, because with 2-ns windows the estimate ($1.54 \pm 0.07\ \mathrm{kcal\ mol^{-1}}$) was still rising at their end; the agent reported the 5-ns estimate with and without the transient and left the choice to the scientist, who excluded the first 1.25 ns. Discarding any prefix between 0.75 and 2.5 ns yields 1.59–1.61 $\mathrm{kcal\ mol^{-1}}$. The agent also cautioned that sequential forward-sweep window seeding cannot rule out hysteresis and that slow $\psi$ transitions (up to ~1 ns at $\phi = 130$–$140°$) exceed the 375 ps bootstrap block length. For steered MD, the work spread was $0.31 \pm 0.62\ \mathrm{kcal\ mol^{-1}}$ ($1.04\, k_{\mathrm{B}}T$, effective sample size 11.8/20), with Jarzynski ($+0.04 \pm 0.11$) and second-order cumulant ($-0.03$) matching umbrella WHAM ($-0.01\ \mathrm{kcal\ mol^{-1}}$) for $F(60°) - F(-110°)$.

**TYK2 RBFE campaigns (complementing Fig. 5 and Supplementary Table 7).** In the **reference network**, the preparation agent inspected all 22 published mappings, noted element-swapping edges (`ejm_50~ejm_55` and `ejm_55~ejm_54`), preserved the 13 input histidine tautomer states (noting automatic PROPKA assignment would flip seven HIE to HID), and tabulated protocol differences against the literature. In the **blind agent-planned network**, a test designed by the scientist (receptor and posed ligands under anonymous names, a budget of about 1,000 GPU-hours, no lookup of the compounds), the agent tested and rejected a ring-size ladder via `map_ligands`, pruned a minimally redundant Kartograf graph from 30 to 24 edges while preserving all low-scoring bridges to peripheral ligands, chose 3 repeats/leg, and flagged missing crystal waters and cyclopropyl stereochemistry for medicinal chemists. A second agent given the same brief independently converged on the same 30-edge planner, mapper, 3 repeats, 11 windows and ring-ladder rejection (not run). In analysis, all 14 shared edges agreed within 0.38 $\mathrm{kcal\ mol^{-1}}$ and per-ligand $\Delta G$ values agreed within 0.28 $\mathrm{kcal\ mol^{-1}}$ r.m.s. Both networks identified `ejm_49` and `ejm_55` as common force-field/binding-mode outliers (consistent with Baumann et al.[12]; excluding the 8 edges through `ejm_55` drops the agent network's $\Delta\Delta G$ RMSE from 0.94 to 0.78 $\mathrm{kcal\ mol^{-1}}$), and flagged the `ejm_54` edges to the scientist as the first to extend, where repeat spread exceeded the per-repeat MBAR estimate (`ejm_55~ejm_54`: 0.29 vs 0.23 $\mathrm{kcal\ mol^{-1}}$; `lig08~lig11`: 0.60 vs 0.31 $\mathrm{kcal\ mol^{-1}}$).

#### Supplementary Video 1 | Building a membrane-protein system via natural language

A scientist works in Claude Desktop, connected over MCP to a live Molpylot session (Molpylot window on the left, conversation on the right). In three requests the language model fetches the KcsA potassium channel (PDB 1BL8) and prepares it for a membrane simulation (pH 7.0, capped termini, restrained minimization); inspects the membrane orientation and the cardiolipins available and embeds the channel in a POPE:POPG:cardiolipin (POPOCL) bilayer (70:20:10) with 0.15 M KCl; and hides the prepared protein so that only the membrane system remains in the 3D view. Every tool call acts on the session shown in the Molpylot window.

#### Supplementary Video 2 | Interactive plots linked to the 3D structure

Three molecular dynamics replicas of ubiquitin analyzed in the Molpylot interface. The radius of gyration and the backbone RMSD are chosen from the grouped catalog of analyses; the RMSD plot shows the replicas one after another or overlaid, with a moving average and the distribution of values, and its points are colored by the radius of gyration. Clicking a time on the plot shows every replica at that frame in the 3D view; clicking a point of one replica shows that replica alone at that frame, and all replicas are then shown again. Secondary structure (DSSP) and per-residue RMSF follow: clicking the most mobile residue centers it in the 3D view, the structure is colored by RMSF, and the helix fraction is overlaid on a second $y$ axis. Finally, the plot is styled for a single journal column (89 mm) and exported as a CSV table and as a figure, which is then shown.

#### Supplementary Video 3 | Conformational landscapes, frame selection and coordinated residues

The same three ubiquitin replicas. A principal component analysis of the C$\alpha$ atoms is shown beside the structure: clicking a point shows that frame, its replica alone, in the 3D view. The projection is then shown as a population map ($-kT \ln P$) and as a scree plot, and the frames of the dense basin are selected as a new entry, whose trajectory is exported. K-means clustering into four clusters follows: clicking a cluster shows its representative structure, and the clusters color the PCA projection. Finally, a distance-fluctuation analysis draws the residue pairs with low fluctuation on the structure, shows the mean distances and the long-range coordinated pairs as a graph, highlights the partners of the most coordinated residue and colors the structure by the local fluctuation.

#### Supplementary Note 1 | Security model for MCP tool execution

Molpylot's MCP server enforces layered operational boundaries around model-driven tool calls:

- **Filesystem isolation.** MCP clients may read and write only inside the server's configured base directory, a private owner-only temporary directory and explicitly user-authorized paths. Without an explicit base directory, the stand-alone server refuses to launch in a filesystem root, home folder, installation folder or read-only directory.
- **Transport authentication.** Embedded loopback HTTP connections require an owner-only locally generated bearer token, a loopback `Host` header (`127.0.0.1` or `localhost`) and no browser `Origin` header unless explicitly allowed; non-conforming requests are rejected prior to dispatch.
- **No model-generated code and strict schemas.** No MCP tool executes arbitrary code supplied by a language model. Calls carrying undeclared arguments are rejected before execution. Session-mutating calls execute atomically inside the undo history (40 steps by default), and dedicated deletion tools carry the MCP `destructiveHint` annotation so clients may prompt for confirmation. Optional server-side call logging records raw arguments, return payloads and durations independently of the model's summary.
- **Indirect prompt-injection mitigation.** Twenty-two tools that return untrusted external text (structure headers, ligand titles, PubMed/UniProt/ChEMBL records, BLAST/structure-search descriptions and job logs) sanitize strings (stripping control characters, ANSI escapes, zero-width/bidirectional/tag Unicode characters and excess length), label them in a structured `external_text` field (the fields and their sources), marked as data-only in server instructions, and scan normalized text against fixed English regular expressions for instruction-like patterns (e.g. prompt overrides, role markers, synthetic tool calls, embedded URLs/HTML or long encoded payloads). Flagged text is preserved for human inspection with an explicit warning in the tool message; entry names taken from molecule titles are restricted to ≤ 64 alphanumeric/basic punctuation characters.
- **Residual boundaries.** Pattern-based marking reduces but cannot eliminate indirect prompt injection (e.g. novel or non-English phrasing, or untagged identifiers re-surfaced through downstream session state), and client-declared model identifiers in provenance records are not cryptographically attested. Structural invariants (no arbitrary code execution, filesystem sandboxing and undo of session changes) remain enforced regardless of text content.

#### Supplementary Note 2 | Agent confinement, standing instructions and representative prompt

The scientist delegated the preparation and analysis of all 18 validation systems to Claude Code agents running Claude Opus 5.5 (medium effort) and supervised their work; the agents were restricted strictly to the stand-alone `molpylot-mcp` server over `stdio`. Shell execution, filesystem access, web tools, sub-agents and all other MCP servers were disabled at the client permission boundary (`.claude/settings.json` and `.mcp.json` of the preparation sessions below; the first line of each block names the file):

```
// .claude/settings.json
{
  "enabledMcpjsonServers": ["molpylot"],
  "permissions": {
    "allow": ["mcp__molpylot"],
    "deny": [
      "Bash", "PowerShell", "Read", "Edit", "Write", "NotebookEdit", "Glob", "Grep",
      "WebFetch", "WebSearch", "Agent", "Skill", "Workflow", "Artifact",
      "mcp__claude-in-chrome", "mcp__visualize", "mcp__terminal"
    ]
  }
}
```

```
// .mcp.json
{
  "mcpServers": {
    "molpylot": {
      "type": "stdio",
      "command": "wsl.exe",
      "args": [
        "-d", "Ubuntu-24.04", "-e", "env", "MOLPYLOT_MCP_MODEL=claude-opus-5-5",
        "/path/to/molpylot-env/bin/molpylot-mcp",
        "--base-dir", "/path/to/validation_md"
      ]
    }
  }
}
```

**Summary of standing instructions (`CLAUDE.md`).** Preparation agents were instructed to: (1) operate exclusively through `mcp__molpylot__*` tools discovered via `find_tools` and `search_docs`; (2) use Molpylot's defaults unless the task asked for something else, or a physical check or the structure required it, documenting every departure with its rationale; (3) resolve all error-level physical checks and explicitly decide and justify every warning; (4) export self-contained SLURM job folders (`write_job`) and one `.msproj` session file per system without running simulations locally; (5) pause and report rather than guess whenever a required operation lacked tool support; and (6) conclude with a structured summary table and a watch-list of items for the scientist to verify before submission.

**Representative open biological brief (rhodopsin `1U19`).** An excerpt of the scientist's brief: the task paragraph, without the standing rules before it and the request for a summary table after it, with the output paths elided ([...]) and the lists run together. It states biological requirements and no force-field, protonation, padding or equilibration directives:

> Prepare bovine rhodopsin in a membrane for 3 independent replicas of 200 ns of production, and write the job (write_job, header "leonardo") in [...]; save the session as [...]. Do not run simulations here.
> The system:
>
> - PDB 1U19, one rhodopsin molecule (chain A), with what belongs to the protein in the native membrane: the retinal bound to Lys296 as a protonated Schiff base, the N-glycans on Asn2 and Asn15, the palmitoylated Cys322 and Cys323, and the acetylated N-terminus. What comes from the crystallization should not be in the simulated system.

- A membrane like that of the rod outer-segment disc, asymmetric: intradiscal (luminal) leaflet: about 60 % phosphatidylcholine, 25 % phosphatidylethanolamine, 15 % cholesterol; cytoplasmic leaflet: about 45 % phosphatidylethanolamine, 25 % phosphatidylserine, 20 % phosphatidylcholine, 10 % cholesterol; acyl chains partly palmitoyl-oleoyl and partly stearoyl-docosahexaenoyl (18:0/22:6), if the lipids available have them.
- Take care of the orientation: the N-terminus and the glycans face the intradiscal side, the C-terminus and the palmitoyl chains the cytoplasmic side. Check which leaflet of the builder ends up on which side of the oriented protein, and that each composition is on the right side.
- Water and salt as for a physiological system.

If something cannot be done with the tools, stop and say why.
